# Bilateral Alignment of Receptive Fields in the Olfactory Cortex

**DOI:** 10.1101/2020.02.24.960922

**Authors:** Julien Grimaud, William Dorrell, Cengiz Pehlevan, Venkatesh Murthy

## Abstract

Each olfactory cortical hemisphere receives ipsilateral odor information directly from the olfactory bulb and contralateral information indirectly from the other cortical hemisphere. Since neural projections to the olfactory cortex are disordered and non-topographic, spatial information cannot be used to align projections from the two sides like in the visual cortex. Therefore, how bilateral information is integrated in individual cortical neurons is unknown. We have found, in mice, that the odor responses of individual neurons to selective stimulation of each of the two nostrils are highly matched, such that odor identity decoding optimized with information arriving from one nostril transfers very well to the other side. Remarkably, these aligned responses are nevertheless asymmetric enough to allow decoding of stimulus laterality. Computational analysis shows that such matched odor tuning is incompatible with purely random connections but is explained readily by Hebbian plasticity structuring bilateral connectivity. Our data reveal that despite the distributed and fragmented sensory representation in the olfactory cortex, odor information across the two hemispheres is highly coordinated.

## Introduction

In senses such as vision and audition, information from paired sensors (eyes or ears) is combined in the brain to perform specific computations that are behaviorally relevant – for example, estimating distance to or the direction of the source of the stimuli (Cumming and DeAngelis, 2001; Grothe and Pecka, 2014). The existence of two sensors for olfaction has prompted hypotheses about their function (Catania, 2013; Esquivelzeta Rabell et al., 2017; Mainland et al., 2002; Porter et al., 2005; Rajan et al., 2006). Information from the two nostrils can be differentiated, or integrated, depending on the goal of such computation for the animal – for example, odor source localization when smooth gradients exist (Álvarez-Salvado et al., 2018; Baker et al., 2018; Catania, 2013; Esquivelzeta Rabell et al., 2017; Gire et al., 2016), or identify relevant smells independent of the odor source location or nostril stimulated (Bojsen-Moller and Fahrenkrug, 1971; Dalal et al., 2020; Eccles, 2000; Kikuta et al., 2008; Kucharski and Hall, 1987, 1988; Mainland et al., 2002). These computations will dictate the nature of the inter-hemispheric communication and the mechanisms of integration of bilateral olfactory cues.

Odors are detected by olfactory sensory neurons in the nose, which project to the ipsilateral olfactory bulb (OB). The OB sends information to multiple ipsilateral brain regions, including the anterior olfactory nucleus (AON) and the anterior and posterior piriform cortices (APC and PPC, respectively) (Giessel and Datta, 2014; Mori et al., 1999; Nagayama et al., 2014; Wilson and Sullivan, 2011) (Figure 1A) In many species, including rodents, the two nares are separated by a septum that minimizes transfer of inhaled odors between nostrils (Eccles, 2000; Kikuta et al., 2008; Wilson and Sullivan, 1999). Therefore, information transfer across hemispheres is largely neural, and originates in the cortex (Bekkers and Suzuki, 2013; Boyd et al., 2012; Haberly and Price, 1978; Kikuta et al., 2008; Scott et al., 1980; Wilson and Sullivan, 2011) (Figure 1A).

**Figure 1:**
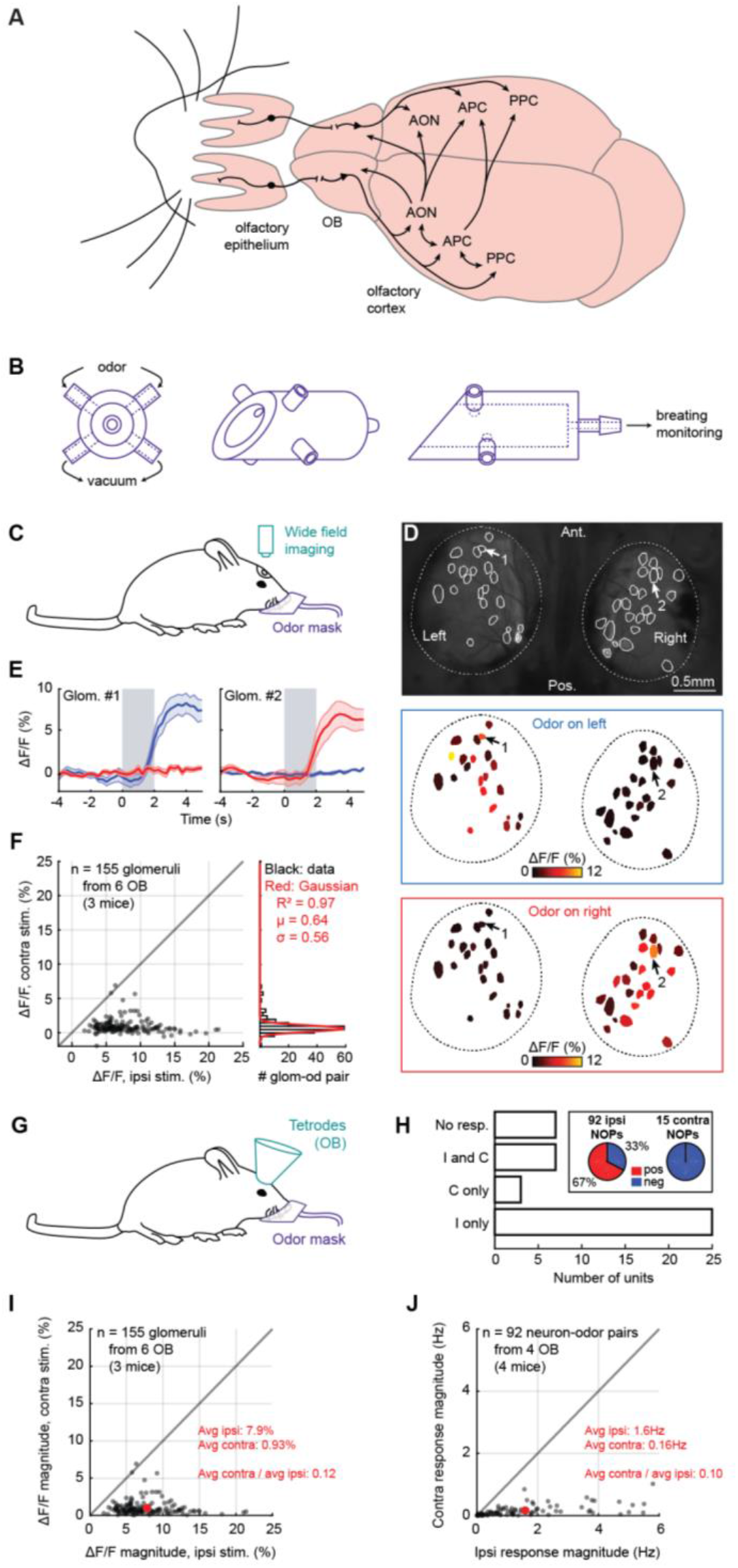
A New Method for Unilateral Odor Delivery. **(A)** Projection between the different olfactory regions. Only the OC (in particular: the AON and APC) sends direct contralateral projections. **(B)** Diagram of the odor mask. From left to right: front, rotated, and side view. **(C)** Calcium imaging setup. **(D)** Exemplar glomerular activity. Example: all glomeruli from OB1, isopropyl tiglate. Arrows: see below. **(E)** Exemplar glomerular activity over time. Grey: odor on (here: isopropyl tiglate). Blue: odor on the left nostril. Red: odor on the right nostril. Average ± SEM. Glomeruli 1 and 2 correspond to arrows 1 and 2 in panel 1D. **(F)** Glomerular responses. Each dot is the response of one glomerulus to one odor (glom-od pair). Histogram: contralateral responses. Red: Gaussian fit. **(G)** Tetrode recording setup. **(H)** Unilateral response profile of M/T cells. NOP: neuron-odor pair. **(I)** Magnitude of glomerular response. Red: average for each side. This panel presents the magnitude of the responses displayed in panel 1F. **(J)** Magnitude of M/T cell response. Only the ipsilaterally responding NOPs are shown. Red: average for each side.

Bilateral cortical interactions have been investigated in other senses including vision, where the brain forms ordered representations of the world using topographic maps, which allow neural matching through precise point-to-point mapping (Cang and Feldheim, 2013; Gu and Cang, 2016; Luo and Flanagan, 2007). However, the olfactory cortex contains no recognizable topography (Bekkers and Suzuki, 2013; Ghosh et al., 2011; Illig and Haberly, 2003; Miyamichi et al., 2011; Pashkovski et al., 2020; Sosulski et al., 2011; Stettler and Axel, 2009; Wilson and Sullivan, 2011), making point-to-point matching of bilateral information through continuous maps implausible. It is possible that odor representations in the two primary olfactory cortex hemispheres are essentially independent, and downstream areas align them for coherent perception, as implicitly proposed by Schaffer et al 2018.

Framing hypotheses about bilateral olfactory processing requires basic information about how single cortical neurons respond to the same odor presented to the two nostrils. Such information is fundamental for uncovering the computations performed, just as details about binocular matching and disparity in cortical responses were for vision (Blake and Wilson, 2011). Individual neurons could respond to ipsi- and contra-lateral odor stimuli in a congruent or disparate manner. There is some evidence for responses in olfactory cortical areas to contralateral odor stimulation, but there is little information on the alignment of ipsi- and contra-lateral responses (Kikuta et al., 2008, 2010; Wilson, 1997). Recent evidence suggests that mitral cells in the OB may have matched responses through organized feedback projections from the anterior olfactory nucleus (AON) par externa (Grobman et al., 2018; Yan et al., 2008). A cortical neuron may then be able to inherit the matched response from mitral cells (Dalal et al., 2020), but a direct evaluation of cortical matching is lacking

To investigate how the olfactory cortex combines information from the two nostrils, we recorded odor-evoked spiking activity of individual neurons in the OC of awake mice. We observed neuron responses to selective odor stimulation through the contralateral nostril in all cortical regions recorded. A significant fraction of OC neurons showed a strong match of ipsi- and contralateral response profiles. Population activity in OC could be used to accurately decode odor identity, as well as side identity. A computational model of the olfactory system suggested that the bilateral overlap we observed in mice is only possible with non-random bilateral projections. Our results provide fundamental and novel insight that defines how ipsi- and contralateral information streams converge and ultimately enhance the processing capabilities of the brain.

## Results

### A New Method for Unilateral Odor Delivery

A major goal of this study was to compare the bilateral odor receptive fields of OC neurons of awake mice. To do so, we needed a way to reliably deliver odors to one nostril at a time. Previous studies have used two different techniques to achieve such unilateral odor stimulations. Either a piece of tubing was inserted in each nostril (Wilson, 1997), or the two nostrils were separated by a plastic septum (Kikuta et al., 2010, 2008). Since mice have very sensitive muzzles, neither of these approaches is applicable to awake animals.

We developed a new facial mask for unilateral odor delivery in awake mice (Figure 1B). On each side of the mask, close to the nostrils, we used a computer-controlled olfactometer to deliver 15 monomolecular odors, one at the time, to the left or the right nostril (Figures S1A-S1B). The mask was connected to an airflow sensor for respiration monitoring (Figures S1C-S1F). We aligned all neuronal odor-evoked activity to the first sniff after odor onset (see Methods).

We tested the side selectivity of the mask through two methods. First, we simultaneously monitored the glomerular activity at the surface of the OBs of OMP-GCaMP3 mice (Isogai et al., 2011) through calcium imaging, while delivering odors to either nostril (Figure 1C) (n=155 glomeruli, 6 OBs, 3 mice). When odors were presented on the contralateral nostril, glomeruli showed little to no response (Figures 1D-1F).

We also performed extracellular recordings of mitral/tufted cells (M/T) in the OB of awake mice using chronically implanted tetrodes (n=42 single units isolated from 4 mice) (Figure 1G, S1G-S1J, and Table S1). M/T cells hardly responded to contralateral odor presentations: out of the 42 isolated neurons, 32 responded to ipsilateral stimulations, while 10 showed significant contralateral responses (Figure 1H). Furthermore, while the ipsilateral M/T cell responses were largely positive, the few contralateral responses were always negative (Figure 1H).

The ratio of contra to ipsilateral response magnitude elicited by odors presented within the mask was 0.12 in our glomerular imaging data (ΔF/F magnitude, average ± SEM, ipsi: 7.9% ± 0.025%; contra: 0.93% ± 7.0e-3%) (Figure 1I) and 0.10 in our M/T recordings (response magnitude, average ± SEM, ipsi: 1.6Hz ± 0.019Hz; contra: 0.16Hz ± 2.1e-3Hz) (Figure 1J). Therefore, on average, odor responses were 8 to 10 times stronger on the ipsilateral side of the mask, compared to the contralateral side. Previous work has indicated that M/T cell responses to a given odor at a concentration of 0.1% were quite similar to the same odor at a concentration of 1% (Bolding and Franks, 2018, 2017). Since the M/T cell responses we observe are substantially smaller for contralateral presentations, the effective odor concentration in the ipsilateral nostril, when odors are presented contralaterally, is likely to be much less than 10% of the initial concentration from ipsilateral presentations.

Together, these results confirmed that odors, when delivered to one nostril through our custom-built mask, activate sensory neurons in the other nostril at least an order of magnitude less than ipsilateral activation.

### Unilateral Odor Tuning of Different Regions of the Olfactory Cortex

We next used our unilateral odor delivery system to characterize how the OC responds to unilateral odor presentations. To do so, we monitored the activity of the AON and APC through extracellular recordings in awake mice (n=7 mice, 1315 neurons total; AON: 3 mice, 384 neurons; APC: 4 mice, 931 neurons) (Figures 2A, S2, and Table S1).

**Figure 2:**
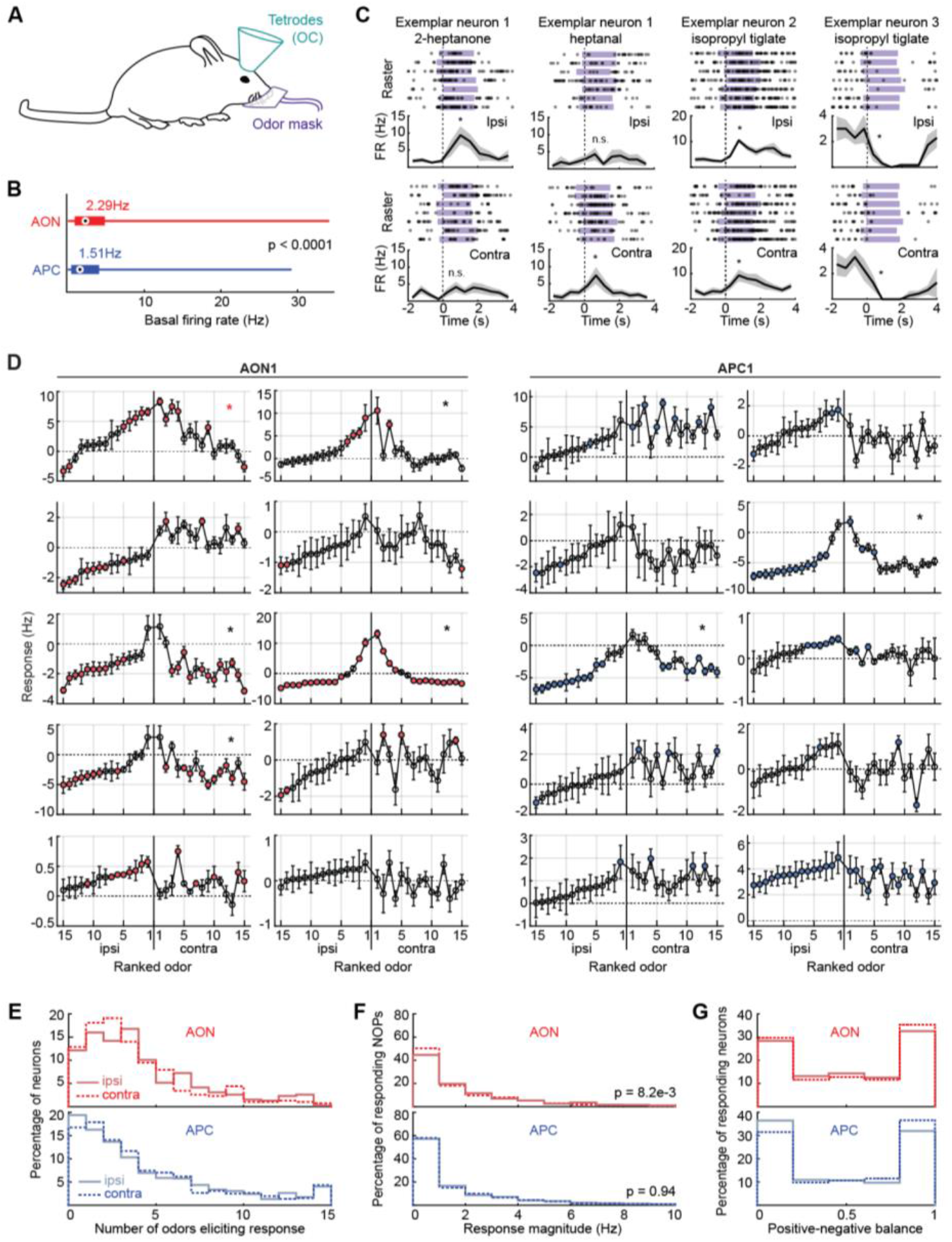
Odor Receptive Fields of Neurons in the Olfactory Cortex. **(A)** Experimental setup. **(B)** Basal activity per region. Circle and value next to it: median. Thick bar: quartiles. Thin bar: range of values. **(C)** Exemplar odor responses over time. Purple: odor delivery. Responses were aligned to the first sniff following odor onset (dotted line). PSTH: average ± SEM. Examples taken from mouse APC1 (see Table S1 for a detailed list of all mice recorded for this study). **(D)** Odor responses, illustration. Neurons randomly picked from mice AON1 (left) and APC1 (right). For each neuron, odors were ranked based on the response magnitude to ipsilateral presentations, which are shown in the left half of each plot. The contralateral responses are then shown with the same rank order of odors for each neuron. Each dot gives the average response ± SEM across the 7 trials, and significant responses are colored red. Asterisk: the ipsi- and contralateral tuning curves of these neurons are visually similar. Further analysis shows that their odor responses are bilaterally correlated. See Figure 3A for details. **(E)** Distribution of the number of odors eliciting a significant response in a given neuron, per region and side. **(F)** Distribution of significant response magnitudes for all neurons and odors, per region and side. **(G)** Positive-negative balance, per region and side. 0 = all negative significant responses; 1 = all positive.

Overall, neurons in the OC had low spontaneous activity (Figure 2B), in accordance with previous studies (Bolding and Franks, 2017; Litaudon et al., 2003; Zhan and Luo, 2010). Spontaneous activity in the AON was significantly higher than in the APC (average spontaneous activity ± SEM, AON: 3.9Hz ± 0.012Hz; APC: 2.8Hz ± 3.6e-3Hz; rank-sum test, p<0.0001).

We found a diversity of responses to ipsi- as well as contralateral odor stimulation in both the AON and APC, including neurons that responded to odors presented to either one or the other nostril, or to both nostrils (Figures 2C-2D). Importantly, in both OC regions, a sizeable fraction of neurons responded to contralateral odor presentations (fraction of neurons significantly responding to at least one odor presented on the ipsilateral nostril, AON: 86%; APC: 81%; contralateral side, AON: 87%; APC: 83%), expanding earlier anesthetized observations to the awake OC (Kikuta et al., 2008, 2010; Wilson, 1997).

OC neurons typically responded to at most a few odors from our odor panel (Figure 2E), as previously reported with bilateral odor stimulations (Bolding and Franks, 2017; Iurilli and Datta, 2017). We found a wide range of response magnitudes across the OC, and contralateral responses were as strong as ipsilateral ones in the APC, and slightly weaker in the AON (Figure 2F; Wilcoxon signed-rank test, ipsi versus contra distribution, AON: p=8.2e-3; APC: p=0.94).

An idiosyncratic feature of olfactory cortical responses is that a neuron that responds to a particular odor with an increase in firing rate, tends to exhibit increases in firing rates for any other odor it responds to; similarly, a neuron having inhibitory response to an odor is also inhibited by any other odor it responds to (Bolding and Franks, 2017; Hu et al., 2017; Kikuta et al., 2008; Otazu et al., 2015; Zhan and Luo, 2010). We find that this feature applies to both cortical regions we examined (Figure 2G).

Overall, we found reliable unilateral odor responses to both the ipsi- and contralateral nostrils in the OC.

### Strong Bilateral Correlations in the Olfactory Cortex

Our data indicate that the OC receives robust contralateral odor information, but it is unclear whether this information is similar to that arriving ipsilaterally. To examine this issue, we related the average odor-evoked responses to ipsilateral against contralateral presentations.

First, we plotted the ipsi- versus contralateral odor responses for each neuron individually. Given the prevalent view of OC connectivity as largely random, we expected OC neurons to show little to no match between their ipsi- and contralateral odor response profiles. To our surprise, we found many neurons with bilaterally-correlated odor responses (Figure 3A) (F-test, significance of the linear model, from left to right neuron, top row: p=0.41; p=0.84; p=0.54; p=0.59; bottom row: p=5.3e-6; p=5.5e-4; p=6.6e-3; p=2.2e-4) (see also Figure 2D, neurons with an asterisk). Overall, for each mouse recorded in the OC, we consistently found a sizeable fraction of bilaterally-correlated neurons (Figure 3B).

**Figure 3:**
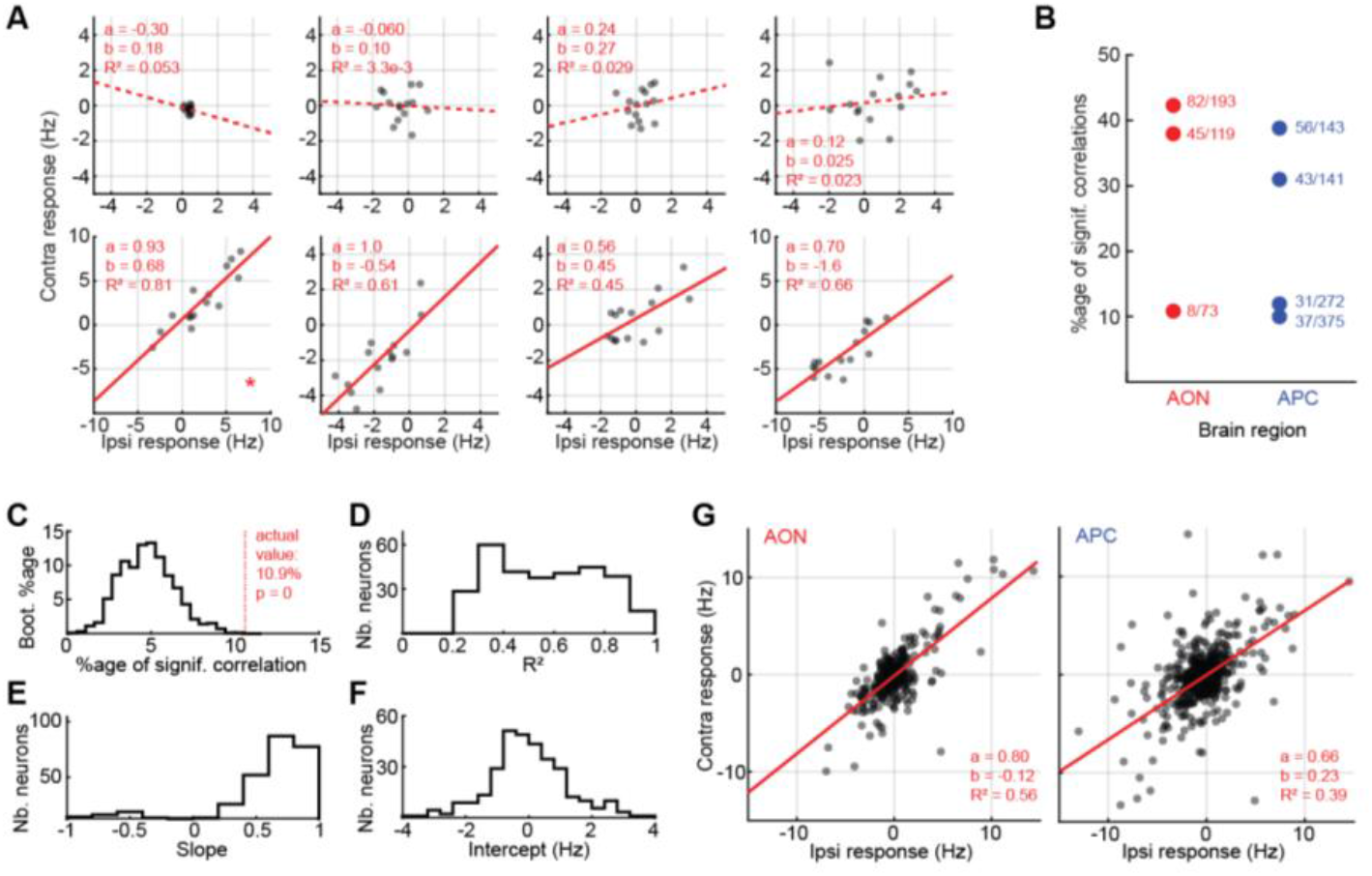
Side-Invariant Odor Responses in the Olfactory Cortex. For panels (A) and (G), red line: linear regression (contra = a × ipsi + b). Solid lines show significant regressions (F-test, p < 5%). **(A)** Exemplar ipsi-versus contralateral responses per neuron. Examples picked from AON1. Each dot is a different odor. The neurons in the bottom panels are bilaterally-correlated. Asterisk: same neuron as in Figure 2D, left panel, red asterisk. **(B)** Percentage of bilaterally-correlated neurons (BCNs) per mouse. Each dot is a mouse. Fractions next to each dot: number of BCNs / total number of neurons. Full circle: the fraction is significantly higher than chance. **(C)** Significance of the percentages of BCNs. Here we show the distribution of chance percentages for AON1, obtained with a bootstrapping procedure. **(D)** to **(F)** Coefficients of determination, slopes, and intercepts of the BCNs in the AON and APC (Nb: number). **(G)** Ipsi-versus contralateral responses per region for odor 1 (isopropyl tiglate). Each dot represents one neuron. For the equivalent graphs across all odors, see Figure S4.

We used a bootstrap strategy to determine that the fraction of bilaterally-correlated neurons in each mouse was significantly higher than what one would expect from chance. In brief, for each mouse, each neuron, we randomly shuffled odor identities on each nostril separately. We repeated this shuffling for all neurons and calculated the resulting percentage of bilaterally-correlated neurons. We reiterated this procedure 10,000 times in order to build a “chance” distribution. Finally, we compared this distribution with the actual fraction of bilaterally-correlated neurons. In all mice recorded in the OC, the fraction of bilaterally-correlated neurons was greater than one would expect from chance (Figures 3B, 3C, and S3A-S3D).

While our initial estimates of the fraction of bilaterally-correlated neurons include all neurons recorded (Figures 3B, S3A), we verified that our conclusions remained true when selecting only the significantly responding (Figures S3B-S3C).

We performed additional analysis to determine the extent to which response variability masks response correlations. We generated individual trials from a Poisson process with rates determined from our recordings. We then enquired how correlated responses would be for two independent sets of 7 trials generated with the same set of data. A large number of such sampling revealed that the fraction of (simulated) neurons that show significant response correlations was on the order of 40-80% even when the ideal response correlations should be 100% (Figure S3D). This analysis indicates that the fraction of significantly correlated neurons in the AON observed in our recordings may underestimate the real fraction of bilaterally-correlated neurons.

Overall, the bilateral correlations were strong (Figure 3D), with mostly positive slopes (Figure 3E), and a distribution of intercepts centered on zero (Figure 3F). In other words, bilaterally-correlated neurons appear to respond to unilateral odor presentations in a mostly side-invariant way, with their ipsi- and contralateral odor responses similar in sign and amplitude.

We wondered if the bilaterally matched responses occurred preferentially in a particular neuron type. We used a classic method for identifying putative excitatory and inhibitory neurons – the width of extracellular action potentials (Frank et al., 2001; Gur et al., 1999; Hassani et al., 2009; Povysheva et al., 2006; Suzuki and Smith, 1985; Swadlow, 2003; Weir et al., 2015). Most of the bilaterally-correlated neurons in the AON and APC had broader spikes and lower spontaneous activity (Figure S3E-S3H), suggesting they are excitatory.

When testing our mask for cross-contamination (Figure 1) we estimated that at most 10% of odors puffed on one side of the mask could leak to the other side. Can this cross-contamination alone explain the side-invariant responses we observed in the OC? To test this hypothesis, we first determined how much each odor from our panel needed to be diluted in the solvent to elicit a PID signal 10 times weaker than our initial dilution (5% volume/volume) (Figure S3I). We then recorded individual neurons in the AON of awake mice (n = 2 mice, 120 neurons) while delivering odors on either side of the mask, both at initial (5%) and new dilutions (i.e. dilutions resulting in PID signals 10 times weaker) (Figure S3J, Table 1). On average, for a given side of the mask, new dilutions elicited neuronal responses 6 to 7 times weaker than our initial dilutions (neuronal response, mean ± SEM, for ipsi presentations, initial dilution: 2.1Hz ± 0.023Hz, final dilution: 0.34Hz ± 2.7e-3Hz; for contra presentations, initial dilution: 2.5Hz ± 0.031Hz, final dilution: 0.37Hz ± 2.5e-3Hz), and the correlation between the two responses was significant (linear regression, ipsi presentations: R^2^=0.18, p<0.0001; contra presentations: R^2^=0.12, p<0.0001). In other words, contralateral cross-contamination in the mask, if any, is not sufficient to explain the presence of side-invariant odor responses.

Finally, we compared the ipsilateral and contralateral population representation in the two OC regions recorded for each odor (Figures 3G and S4A-S4C). Consistent with our analysis of individual bilaterally-correlated neurons, we found that unilateral population representations were significantly and strongly correlated for all odors tested in the AON and APC (Figures S4A-S4B) (see Table S2 for detailed statistics). Analyzing population correlations for each odor individually ensures that the correlations we observe are not driven by few neurons responding strongly and similarly to all odors, which would otherwise create an artificially strong positive correlation. Furthermore, a bootstrap analysis similar to what we described above confirmed that strong correlation we observed for each odor individually were unlikely to occur by chance (Figures S4C-S4D). When considering each mouse separately, our conclusions held true for most odors (Figure S4E).

Together, these data indicate that the odor information arriving contralaterally to the OC strongly matches that arriving ipsilaterally.

### Accurate Decoding of Odor Identity from Contralateral Activity

Odor selectivity in the OC allows a population representation of odor identity, and odor identity can be decoded from the activity of a sufficient number of neurons (Bolding and Franks, 2017). If odor responses are similar for ipsi- and contralateral presentations, decoding of identity should be transferrable. To formally test this, we asked a linear classifier to decode odor identity from OC responses to stimuli in one nostril, after we trained it with OC responses to stimuli in the opposite nostril.

In brief, at each instance we created a training set by randomly removing one trial per odor (Bolding and Franks, 2017). If we were testing on the same side as we trained (e.g. train ipsilaterally, test ipsilaterally), then we tested the classifier on the trials outside of the training set. If we were testing on the opposite side (e.g. train ipsilaterally, test contralaterally), then one trial per odor from the opposite side was randomly chosen to form the test set (Figure 4A). We performed all our decoding analyses over an odor response window of 2s, as it maximized performance of odor decoding on the same side (Figure S5A).

**Figure 4:**
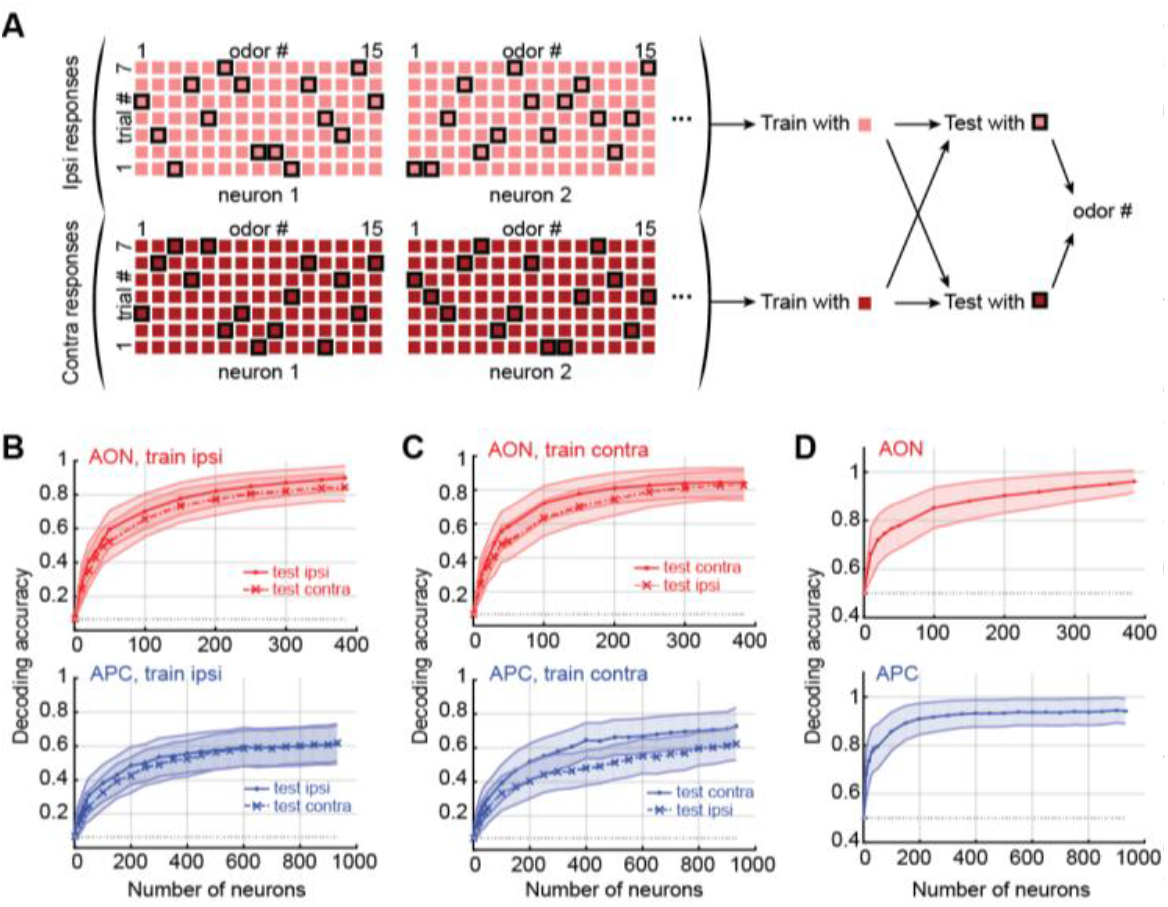
Decoding Odor Identity and Laterality. **(A)** Diagram of the odor decoding process. **(B)** and **(C)** Odor identity decoding per region. Mean ± SEM over 200 repetitions of the decoding process. For (B), the decoder was trained with data from ipsilateral presentations. For (C), the decoder was trained on contralateral odor responses. **(D)** Side identity decoding per region. Mean ± SEM over 200 repetitions of the decoding process. For (B) to (D), grey dotted line: chance.

In both the AON and APC, the decoder performed with similarly high levels of accuracy when tested on data from the same or the opposite side (Figures 4B-4C). In other words, odor identity decoding optimized with responses to stimulation from one nostril transfers very well to stimulation of the other side.

Note that the seeming decrease of decoding performance from AON to APC is most probably due to variability across mice (Figure S5B). More precisely, the three mice with overall weaker and fewer odor-evoked responses (i.e. mice AON2, APC1, and APC2) showed lower odor decoding performances, no matter the side tested (Figure S6). In addition, AON2 is the mouse from which we isolated the fewest neurons (73 neurons, while all other OC recorded mice have over a hundred neurons each) (Table S1). Predictably, in these same three mice, the fractions of bilaterally-correlated neurons, while significantly higher than chance, were the lowest.

Rodents are able to determine the direction of the odor stimulation at short distances (Esquivelzeta Rabell et al., 2017; Rajan et al., 2006). We asked whether population activity in a single hemisphere differs sufficiently for stimulation of ipsi- and contralateral nostril to allow decoding of intensity differences between the two sides. Side identity, or laterality, could indeed be decoded with high accuracy (Figure 4D), which confirms that odor representations from both sides, despite similarities, remain distinct and separable.

Together, our data indicate that odor decoding is transferrable across sides. Ipsi- and contralateral odor representations, while distinct, share a high degree of similarity.

### A Computational Model of Bilateral Matching in the Olfactory Cortex

How is one OC responding so similarly to inputs from two distinct routes/nostrils? So similarly, in fact, that a linear classifier trained on one set of representations can decode the other?

An appealingly simple resolution might suggest itself based on recent theoretical work. It has been shown that if two correlated representations are projected through the same random matrix the output representations are also correlated (Babadi and Sompolinsky, 2014; Schaffer et al., 2018). Thus, one could hypothesize that any interhemispheric projection between two OCs could preserve the structure inherited from the underlying OB representations, hence explaining the similarity. However, this argument is insufficient, as illustrated in Figure 5. The correlations in glomerular neural representations of the 15 odors used in our study are shown in Figure 5B. These correlations are partially preserved when projected through two different random matrices (to represent the OB to OC projections of each hemisphere) (Figure 5C, upper and lower block diagonals). However, the two OC hemispheres connected with random projections will not yield correlated responses to the same odor presented to the left and right side (Figure 5C). In the supplementary material we show that, even in the limiting case where the response of ipsi- and contralateral cortices are identical, unstructured interhemispheric connectivity ensures that the responses of one cortex to odors presented in different nostrils will be uncorrelated. Hence, there must exist some structured connectivity that is aligning the cortical responses (Figure 5D; note the much higher similarity values in the two off-diagonal blocks).

**Figure 5:**
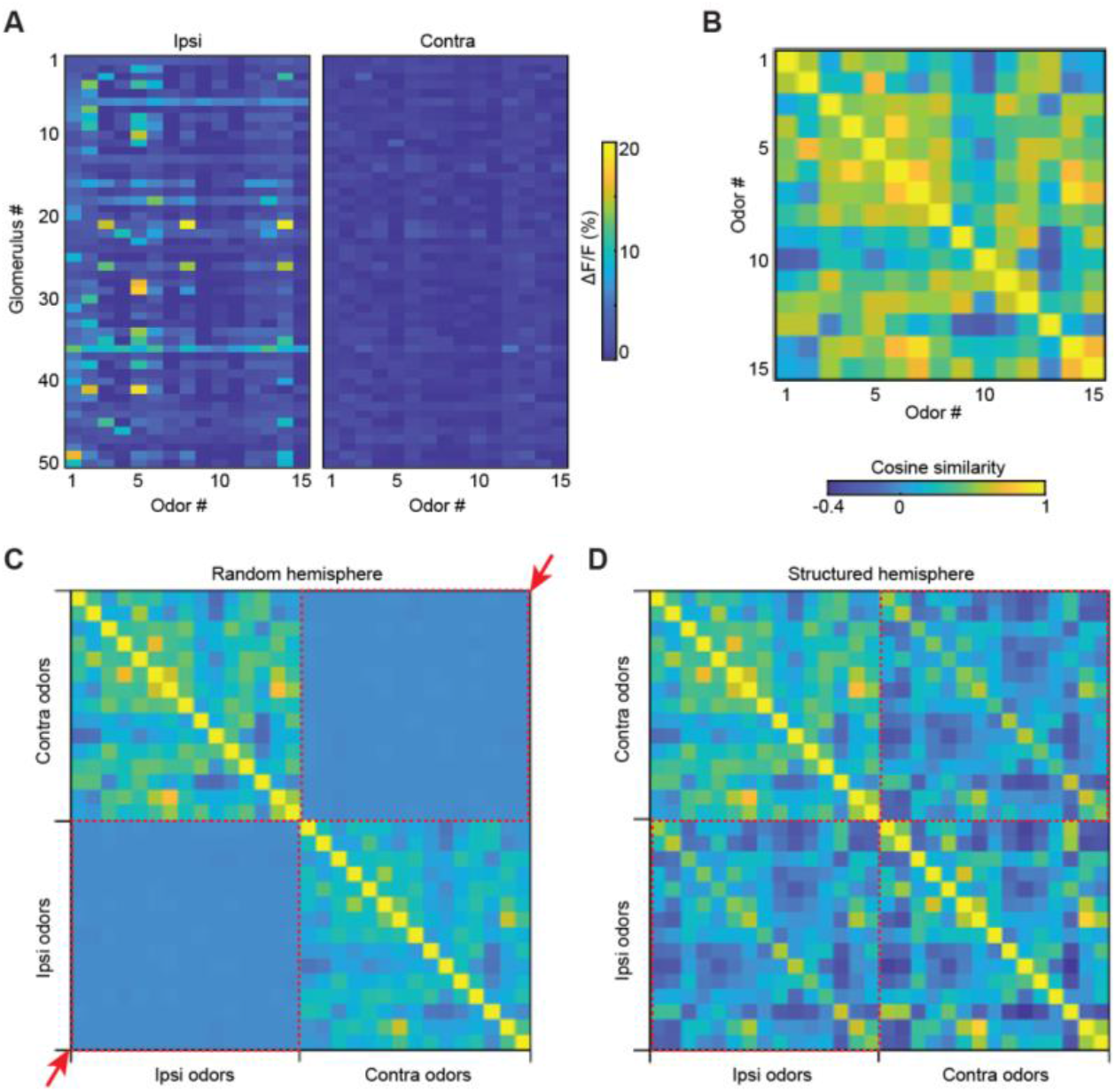
Structure is necessary for alignment between responses to different nostrils. **(A)** Odor tuning of 50 randomly selected glomeruli. **(B)** The OB odor-odor cosine similarity matrix measured using calcium imaging. Color scale at right applies to panels (B) to (D). **(C)** We modeled the two OCs and connected them using a random matrix (see methods), the correlations between odors from the same nostril are preserved, but correlations between nostrils are completely lost (quadrants pointed to by red arrows and red dotted squares). **(D)** Including a structured cross-cortical connectivity produces correlations between ipsi- and contralateral odors.

We then asked what type of structured interhemispheric connectivity is sufficient to achieve the observed tight alignment of responses from different nostrils. Through computational modelling we tested a simple and plausible model for structured matching, a Hebbian connectivity matrix that links together the same-odor representations in each hemisphere. We combined this Hebbian matrix, (*G_Structured_*), with a random matrix, (*G_Random_*), which models unstructured elements of connectivity and whose elements are drawn independently from a Gaussian distribution. We vary the degree of structure using a parameter *α*. Explicitly:

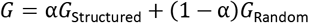

The parameter *α* took values between 0 and 1 with *α* = 0 corresponding to purely random connectivity and *α* = 1 to purely structured connectivity. The random and Hebbian components were scaled appropriately such that *α* represents the proportion of contralateral input magnitude that is formed by structured connectivity.

Our modeling strategy began by creating a cortical representation of odors presented in the ipsilateral nostril, based on the AON. Random OB-to-OC connectivity indicates that each cortical olfactory neuron samples a random subset of glomeruli. This implies that the input to each cortical neuron is statistically independent, and that the response of one cortical neuron to different odorants can be modelled as a thresholded multivariate gaussian whose covariance matrix can be extracted from activities in the OB (Figure 6A, see Methods). We validated this model of a single AON against neuronal recordings: our model replicated the population statistics remarkably well with minimal parameter adjustment (Figures 6B-6D; Figures S7A-S7C).

**Figure 6:**
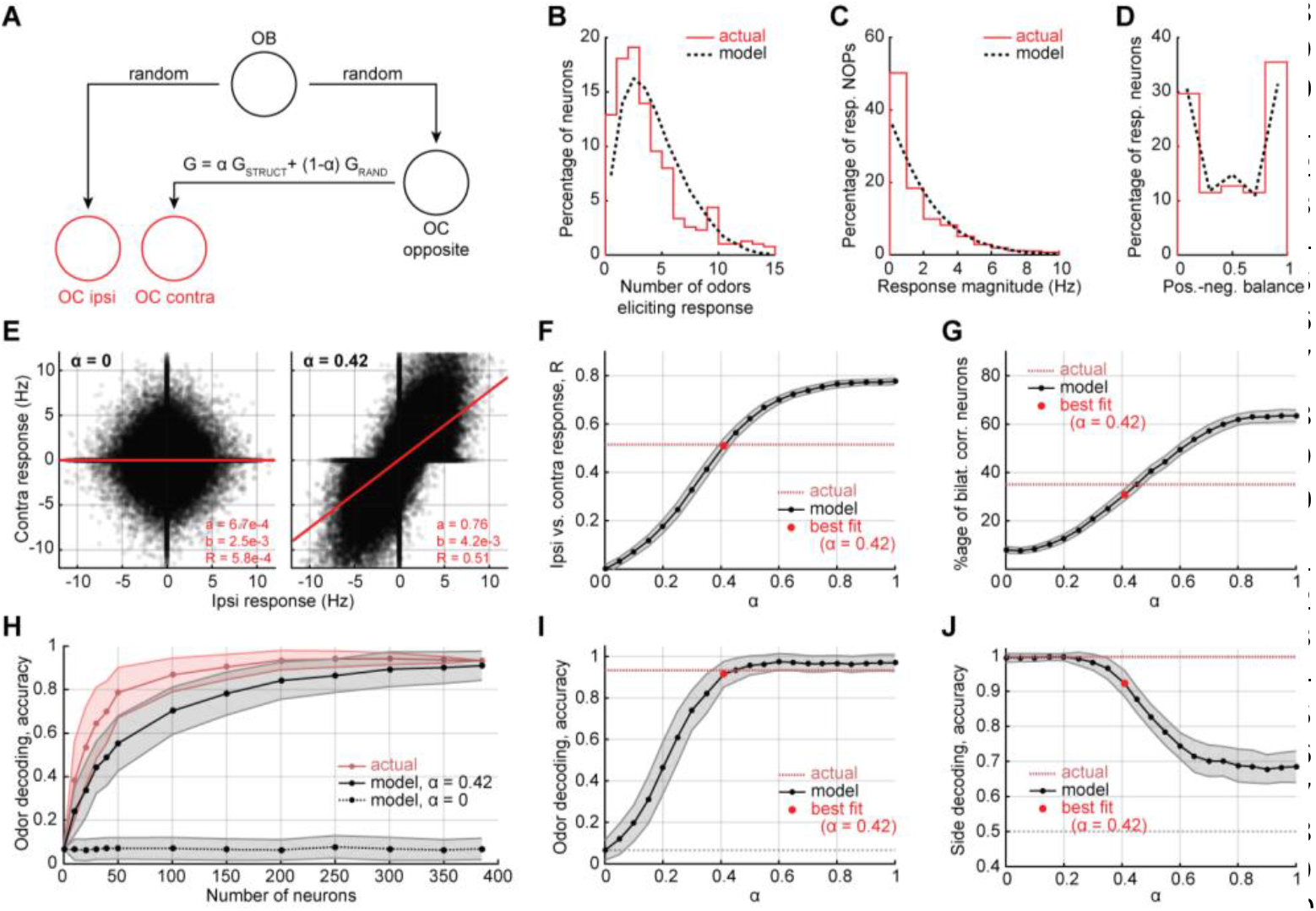
Modeling Olfactory Cross Cortical Connections. **(A)** Diagram of the model. **(B)** to **(D)** Model validation. Contralateral odor responses from the model are overlaid with actual contralateral AON responses (Figures 2E-2G). **(E)** to **(J)** Various measurements of cross-cortical alignment from the model. **(E)** Contra- vs ipsilateral responses for random (alpha = 0) and semi-structured (alpha = 0.42) projections. **(F)** Contra- vs ipsilateral response correlations R for various alpha values. **(G)** Percentage of bilaterally-correlated neurons for various alpha values. **(H)** Odor decoding accuracy for random (alpha = 0) and semi-structured (alpha = 0.42) projections. Red: same as Figure 3B, AON, “test contra”. **(I)** Odor decoding accuracy for various alpha values. **(J)** Side decoding accuracy for various alpha values. For (E) to (H), measurements from the model were calculated by sampling 385 neurons 200 times. The best fit alpha (0.42) was found by comparing to AON data (n = 385 neurons from 3 mice).

We then linked two simulated AONs using the connectivity matrix *G*. The computational analogue of the cortical representations of an odor presented contralaterally is the representation after mapping through the interhemispheric matrix, *G*. Thus, by comparing the simulated and measured alignment of ipsi- and contralaterally presented odors as we vary α, we can ask if this simple Hebbian structure is sufficient to recapture the degree of alignment observed.

We compared the alignment using four metrics: the overall correlation of odor population representation across sides (Figures 6E-6F; similar to Figure 3G), the fraction of bilaterally-correlated neurons (Figure 6G; similar to Figure 3B), the accuracy of odor identity decoding (Figures 6H-6I and S7D; similar to Figures 4B-4C), and the accuracy of side identity decoding (Figure 6J; similar to Figure 4D).

The four computed measurements best matched our actual AON neuronal recordings with a non-zero *α* (approximately 0.4). This result did not vary when we changed the size of the modeled OC population (Figures S7E-S7H). This suggests that a significantly Hebbian connectivity between cortices is sufficient to explain the observations.

We chose this particular structured connectivity, the 15 odor Hebbian matrix *G_Structured_*, for its appealing simplicity and biological plausibility. For example, early in life the animal might have smelt these odors in both nostrils and associatively matched the responses between hemispheres. However, there are many other sufficient alignment schemes that do not rely on having experienced specifically these 15 odors; we implemented two such examples in the supplementary material. The first uses a set of Hebbian matching experiences that span the subspace in which the 15 odorants lie. Roughly 6 of these are needed to achieve similar levels of alignment (Figure S8G-S8J) showing that a smaller number of more general early learning experiences could also explain the alignment seen in the data. The second technique uses Singular Value Decomposition to create a low rank approximation of *G_Structured_*; again, a rank of about 7 is necessary to match the data (Figure S8C-S8F). Effectively, any connectivity that links same-odor representations in each hemisphere together will be able to match the alignment seen in the data.

## Discussion

In many animals, odor information collected by the two nostrils remains separate until it reaches secondary brain areas (Bojsen-Moller and Fahrenkrug, 1971; Eccles, 2000; Kikuta et al., 2008). How the two streams of information are integrated, as well as the functional role of this integration, remain unclear.

Here, we reveal that a sizeable fraction of neurons in the OC are bilaterally-correlated, meaning that they have matched ipsi- and contralateral odor tuning. Functional alignment of responses was also confirmed by the ability to decode odor identity from neural populations trained using only odor responses from the opposite nostril. Finally, our computational model suggests that such bilateral matching requires structured interhemispheric projections. Our findings shed light on how brains make sense of this bilateral experience in the olfactory system.

### Importance of the Integration of Bilateral Information

In many senses, information is collected through a pair of sensors and is integrated within the brain. Bilateral perception allows us to perform vital behavioral computations such as sound localization or depth perception. Since the two nostrils may sample independent pockets of air (Eccles, 2000; Kikuta et al., 2008; Wilson and Sullivan, 1999), can bilateral information be used to discover features of the odor stimulus not readily available to single nostrils? Animals may compare of odor concentration across both nostrils to locate odor sources (Catania, 2013; Esquivelzeta Rabell et al., 2017; Gire et al., 2016; Parthasarathy and Bhalla, 2013), which requires bilateral integration and comparison of different signals. Two brain regions that seem to fulfil this criterion are the AON and APC (Kikuta et al., 2008, 2010; Wilson, 1997). Recent studies have indicated that the AON may be sensitive to the differential activation of left and right nostrils, therefore computing the direction of the stimulus in front of the animal (Esquivelzeta Rabell et al., 2017). Our own data shows that side identity can be decoded from odor responses, which provides further evidence for differential processing (Figure 4D).

In contrast, many behaviors require the animal to make inferences independent of the nostril stimulated. One simple example can be found in our everyday life: one nostril is usually more blocked than the other, and which nostril gets clogged changes several times a day (Eccles, 2000). Nevertheless, we can identify smells and have a unified olfactory experience. In the same fashion, animals need to be able to identify relevant smells like food or a predator, no matter where the odor source is relative to their nose. Indeed, there is evidence for nostril-independent recognition of odors: rats trained to associate odors with reward using only one naris can generalize to stimuli delivered through the other nostril (Kucharski and Hall, 1988, 1987; Yan et al., 2008). This behavior requires interhemispheric communication, suggesting a convergence of ipsilateral and contralateral representations in higher brain areas (Dalal et al., 2020).

### Robust Contralateral Odor Responses in the Olfactory Cortex

Here, we developed and tested a new method to deliver odor stimuli selectively to individual nostrils in awake, freely breathing mice (Figure 1). We performed several experiments to confirm selective stimulation. First, PID recordings indicated that odors were delivered at the same concentration on either side and are not detectable on the side contralateral to the stimulus. Second, direct measurements of sensory input to the OB indicated that glomerular responses to contralateral stimulation were below detection threshold. It is possible that some low-level cross contamination went undetected by the PID and glomerular recordings, but it is unlikely that such weak inputs could contribute to contralateral responses that are as strong as those to ipsilateral stimulation, as shown by our recordings in AON to 10-fold dilute odors (Figure S3J).

Third, recordings from putative M/T cells in the OB using tetrodes revealed few detectable responses to contralateral nares stimulation. These results seem at odds with a recent report suggesting that topographic projections from contralateral AONpE to M/T cells support bilaterally matched odor responses (Grobman et al., 2018). A possible source of the discrepancy is the much lower concentration of odor used in our mask. However our observations confirm former work from various teams regarding the near absence of contralateral M/T response in the OB (Kikuta et al., 2008, 2010; Wilson, 1997). Our results are also consistent with the known projections from the AON pars externa to the GABAergic granule cells of the contralateral OB (Schoenfeld and Macrides, 1984; Yan et al., 2008), since the infrequent significant responses we observed in M/T cells to contralateral nostril stimulation were inhibitory. The minimal responses in glomeruli and M/T cells also argues strongly against bilateral return of odors from the respiratory system during exhalation.

### Bilateral Integration in the Olfactory Cortex is not Random

Our first key finding is that responses to the stimulation of the contralateral nostril could readily be detected in the AON and APC (Figure 2). This is not entirely surprising, since crossed projections already occur at the level of the AON and APC. Previous studies indicated that contralateral stimulation evoked fewer and weaker responses in the AON (Kikuta et al., 2008). However, these studies recorded from a small number of neurons in anesthetized animals. While we confirmed the presence of weaker and fewer responses to contralateral stimulations in the AON, the differences were much smaller than previously reported. Another recent study implicated AON in interhemispheric communication using a variety of methods, but the responses of neurons in this area were not measured (Yan et al., 2008). Finally, one previous study reported the presence of neurons responsive to contralateral naris stimulations in the APC of anesthetized rats, but did not test the correlation of the responses between the two nares (Wilson, 1997).

We found that the odor information arriving ipsilaterally to the OC strongly matches that arriving contralaterally (Figure 3). Our control experiments indicated that less than 10% of an odor delivered to one nostril may leak to the other nostril, and we found that this potential 10% leak is not sufficient to explain the presence of bilateral neurons. Furthermore, our data offer strong hints that principal neurons are a major part of the ensemble of neurons that have matched responses to stimuli from both nostrils (Figure S3). This feature would allow downstream areas receiving olfactory cortical inputs to readily have access to side-stable odor responses. Indeed, we showed explicitly that odor decoding is transferrable across sides (Figure 4), as expected from the presence of matched responses in the OC.

Finally, we considered how the prevalent view of OC connectivity could explain our data (Figure 5). Recent studies have shown that two representations projected through the same random matrix remain correlated (Babadi and Sompolinsky, 2014; Schaffer et al., 2018). In addition, another study reported that population response correlations between odors is shaped in the cortex, presumably by experience, such that some response similarities in the OB are enhanced and others are suppressed (Pashkovski et al., 2020). However, these studies do not address alignment due to projections between cortices. Here we implemented a computational model of the olfactory system to test matching of ipsilateral and contralateral odor tuning in the OC as a function of structure in bilateral projections. We showed that, in our model, purely random projections cannot explain our OC results. That is, even if the two hemispheres have the same ordered representations for similar odors, unstructured interhemispheric cortical projections will disrupt these correlations.

### Reconciling Our Results with Known Connectivity Schemes

How can the responses of cortical neurons to selective stimulation of each of the two nostrils be matched? First, there may be topographically matched crossed projection somewhere in the olfactory system allowing bilateral response matching. The only known candidate for such matching is the projection from AON pars externa to OB (Schoenfeld and Macrides, 1984; Yan et al., 2008). This projection, however, should result in net inhibition of M/T cells since its major target are the GABAergic granule cells (Schoenfeld and Macrides, 1984; Yan et al., 2008) (but see Grobman et al., 2018). Furthermore, previous studies (Kikuta et al., 2010, 2008; Wilson, 1997) and our own data (Figure 1) suggest that this projection is too weak to support the widespread bilateral matching we observed in the OC. However, M/T cells that respond to contralateral odor presentation in a correlated way (Grobman et al 2018) may report to ipsilateral OC neurons the same information for ipsi- and contralateral stimulation under some conditions, for example, high odor concentrations. Even so, unstructured interhemispheric cortical projections will disrupt this potential alignment in the cortex. Therefore, additional mechanisms, such as Hebbian plasticity, are needed to explain our observations.

A second way in which matched responses may arise is through fine-scale order in anatomical connectivity that somehow allows the same glomerular output channels from both OBs to converge on individual neurons. Such structure seems implausible and has no precedent but remains formally possible.

A third explanation that seems likely because of its simplicity and local nature is that synaptic connectivity in cortical regions is refined by Hebbian plasticity (Best and Wilson, 2003; Franks and Isaacson, 2005; Johenning et al., 2009) to allow matched responses from both sides. During their normal experience, mice will frequently inhale similar odors through both nostrils. This can allow associative plasticity in cortical areas to select for matched inputs from both hemispheres, similar to, for example, the well-studied map development in the visual system (Bednar and Wilson, 2016; Gu and Cang, 2016; Hubel and Wiesel, 1962). This hypothesis of experience-dependent matching of bilateral inputs to individual cortical neurons is open to experimental testing.

## Methods

### Contact for Reagent and Resource Sharing

Further information and requests for resources and reagents should be directed to and will be fulfilled by the Lead Contact, Venkatesh N. Murthy.

### Experimental Model and Subject Details

Male C57Bl/6J and OMP-GCaMP3 (Isogai et al., 2011) mice were used in this study. All the mice were 3 to 8 months old at the time of the experiments. Mice were singly housed after chronic tetrode implantation. A summary of the mice used for tetrode recording can be found in Table S1. All experiments were performed in accordance with the guidelines set by the National Institutes of Health and approved by the Institutional Animal Care and Use Committee at Harvard University.

### Bilateral Odor Stimulation

The odor panel was composed of 15 monomolecular odors, purchased from Millipore Sigma: isopropyl tiglate, ethyl tiglate, propyl acetate, isoamyl acetate, ethyl valerate, hexanal, heptanal, allyl butyrate, citronellal, hexyl tiglate, 4-allyl anisole, isobutyl propionate, 2-heptanone, ethyl propionate, and eucalyptol (Figure S1A). The odors were diluted in diethyl phthalate, also purchased from Millipore Sigma (final odor dilution: 5% volume/volume). In addition to these 15 odors, we also tested the response to diethyl phthalate alone, referred as the “blank” trials (Methods, “Neuronal Recording” section). The olfactometer was controlled by a custom LabVIEW script (National Instruments).

Odors were delivered through a custom-built facial mask (Figure 1B). The mask was made of a large piece of tubing surrounding the mouse’s muzzle. On each side of the mask, close to each nostril, a piece of tubing delivered odors at a small flow rate (50 standard cubic centimeters per minute). Also on the mask, right below each odor delivery tube, a vacuum ensured proper air cleaning. We confirmed that the mask could deliver similar amounts of odor on each side of the mask with a photoionization detector (miniPID 200B, Aurora Scientific, Canada) (Figure S1B). We confirmed the absence of contralateral stimulations through calcium imaging at the surface of the OB and M/T cell recordings (Figure 1).

Odors were delivered for 2s for each trial. To avoid odor habituation, a period of 15s separated consecutive trials (i.e., 15s from the end of a trial to the beginning of the next trial). Each odor was delivered in two configurations: ipsilateral and contralateral to the implantation. Each odor delivery was repeated 7 times. Therefore, each recording session contained (15 *odors* + 1 *blank trial*) × 2 *sides* × 7 *repetitions* = 224 odor presentations. The order of odor presentations was randomly shuffled for each recording session.

### Breathing Monitoring

The face mask was connected to an airflow sensor (Honeywell AWM3300V, Morris Plains, NJ) to monitor the animal’s respiration during the experiments (Figure 1B) and later align neuronal activity with the sniffs (Figure 2C) (Bolding and Franks, 2017). The accuracy of the face mask in monitoring respiration was tested against intranasal pressure transients (Figures S1C-S1F) as described previously (Reisert et al., 2014).

### Chronic Tetrode Implantation

Mice were anesthetized with an intraperitoneal injection of ketamine and xylazine (100 mg/kg and 10 mg/kg, respectively), then placed in a stereotaxic apparatus. After the skull was cleaned and gently scratched, a custom-made titanium head bar was glued to it. A small craniotomy was performed above the implantation site, before 8 custom-built tetrodes (Chang et al., 2013) were lowered together into the brain (coordinates for the AON pars principalis: antero-posterior 3mm, medio-lateral 1.5mm, dorso-ventral 2.3mm - APC: antero-posterior 1.6mm, medio-lateral −2.8mm, dorso-ventral 3.4mm - PPC: antero-posterior 0.3mm, medio-lateral 3.9mm, dorso-ventral 3.5mm - OB: antero-posterior 1.2 mm, medio-lateral 1.1 mm, dorso-ventral 0.3mm; all antero-posterior and medio-lateral coordinates are given relative to Bregma, except for the OB, for which they are given relative to the junction of inferior cerebral vein and superior sagittal sinus; all dorso-ventral coordinates are given relative to the brain surface). A reference electrode was implanted on the occipital crest. All the mice were implanted in the right hemisphere, except mouse APC2, which was implanted in the left hemisphere. We did not find any difference between APC2 and the other mice. The whole system was stabilized with dental cement. Mice were given a week of recovery before any new manipulation.

### Neuronal Recording

A week after chronic tetrode implantation, mice were progressively habituated to stay calm while head-fixed on the recording setup. The habituation process typically took a week. Brain activity was then recorded once a day, always at the same period of the day, while delivering odors to the ipsilateral or contralateral nostril. The mice were awake, head-restrained, and freely-breathing during all the recordings.

Since the tetrode headstage allowed for the tetrodes to be finely adjusted up or down, the tetrodes were slightly lowered in the brain after each recording session (around 40μm deep), ensuring that different neurons were recorded each day.

Electrical activity was amplified, filtered (0.3-6kHz), and digitized at 30kHz (Intan Technologies, RHD2132, connected to an Open Ephys board). Single units were sorted offline manually using MClust script (Schmitzer-Torbert et al., 2005) written for MATLAB (MathWorks). Units with more than 1% of their inter spike intervals below 2ms refractory period were discarded. Units displaying large changes of amplitude or waveform during the recording were also discarded (Rey et al., 2015) (Figures S1G-S1I). The position of the tetrodes in the brain was confirmed post-mortem through electrolesion (200μA for 4s per channel) (Figure S2).

The average response *R* to an odor *o* (in Hz) over the 7 trials *t* was calculated as followed:

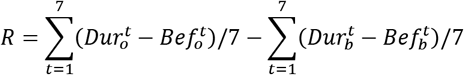

where, for each trial *t*,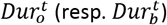 is the firing rate during the odor (resp. “blank”) delivery, while 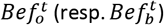 is the firing rate before the odor (resp. “blank”) delivery.

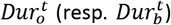 was calculated for the first 2s following the first sniff after odor (resp. “blank”) onset. 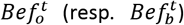 was calculated for the 4s preceding odor (resp. “blank”) onset.

To determine the significance of a response to an odor *o* over the 7 trials *t*, we compared the activity evoked by the odor 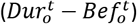 to the “blank” activity 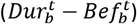 (Wilcoxon signed-rank test, critical value set to 5%) (Bolding and Franks, 2017).

Finally, we investigated whether a neuron that responds to a particular odor with an increase in firing rate, tends to exhibit increases in firing rates for any other odor it responds to; and similarly, whether a neuron having inhibitory response to an odor is also inhibited by any other odor it responds to. To do so, we calculated a “positive-negative balance” metric. For each neuron significantly responding to at least one odor, the positive-negative balance corresponds to the fraction of significant responses that are positive. For example, if a neuron has a positive-negative balance of 0.5, then we know that, among the odors eliciting a significant response, half of them elicit an increase of activity, while the other half elicit a decrease. Similarly, a balance of 0 means that all the significant odor responses are negative, and a balance of 1 indicates that all the significant responses are positive.

### Acute Calcium Imaging

Mice were anesthetized with an intraperitoneal injection of ketamine/xylazine mixture (100 mg/kg and 10 mg/kg, respectively). After the skull was cleaned and gently scratched, a custom-made titanium head bar was glued to it. A large craniotomy was performed above the two OBs and covered with 1% low melting point agarose (Thermo Fisher). While the mouse was still anesthetized, calcium activity was imaged above the craniotomies with a custom-made epifluorescence microscope (10X objective, Olympus, NA 0.3; blue excitation LED, Thorlabs; emission filter, 500–550nm, Thorlabs; CMOS camera, The Imaging Source) at 4 frames/s, while presenting odors with the custom-built facial mask (Figures 1A and 1B).

The acquired sequences of images were converted to ΔF/F images (change in fluorescence over resting fluorescence). Then the maximal response at each pixel was calculated for each of the 15 odors, for each stimulus configuration (right or left nostrils), and these maximum response maps were used to manually identify glomeruli for further analysis. Finally, for each odor, for each stimulus configuration, the response at each glomerulus was quantified by integrating the ΔF/F signal 4s after odor onset (which corresponds to the 2s of odor presentation, and the 2s immediately following).

### Decoding Task

#### Neuronal Decoding

For each neuron we created a feature vector by calculating the summed spike count in a 2s window after the mouse first sniffs the odor and subtracting from it the same sum from the ‘blank’ odor averaged across seven trials.

Odor identity decoding ability was assessed using linear classifiers and leave-out cross validation on a pseudopopulation made up of all neurons recorded in a particular cortical area – AON or APC (Bolding and Franks, 2017). At each instance we chose a particular dataset (either ipsilateral or contralateral odor presentation) and formed from it a training set by randomly removing one trial per odor. We calculated mean responses for each odor by averaging across the training set. We then chose our test set in two related ways. If we were testing on the same side as we trained (e.g., train ipsilaterally, test ipsilaterally), then the 15 trials (one per odor) outside the training set were used as the test set. If we were testing on the opposite side (e.g., train ipsilaterally, test contralaterally), then one trial per odor from the opposite side was randomly chosen to form the test set. Classification accuracy was measured by assigning each test trial to the closest Euclidean mean odor response and calculating the percentage of trials correctly assigned (Bolding and Franks, 2017). This process was repeated 200 times using different random test and train trial choices.

In order to examine the effect of changing number of neurons on classification accuracy, we randomly selected neurons from the pseudopopulation and performed classification using only the selected cells. This process is repeated 200 times. We then report the average accuracy and standard deviation over those 200 selections of neurons and, for each set of neurons, 200 sets of train/test trials in Figures 4 and S5.

A very similar process is employed for side decoding. One trial per side was removed, the remaining trials are averaged to form a mean side response for both ipsilateral and contralateral presentations. The removed trials are then classified to the nearer mean side response and accuracy is the percentage correctly assigned. This is repeated 200 times for different random removed trial choices.

#### Comparisons to the Theoretical Model

Our model predicts the trial-averaged neural responses. In order to make comparisons to the neuronal data, we designed a new classification task. For simulations and trial averaged neuronal data we performed odor decoding as follows: we assigned each of the 15 odor centroids from one side (contralateral or ipsilateral) to their closest neighbor on the other side. This classification was correct if an odor from one side was classified to the same odor on the other side. The accuracy is then the proportion of odors correctly assigned. Again, average accuracy and standard deviation over 200 randomly chosen neurons are reported in Figures 4 and S5. One exception to the procedure above is Figure S7D, right panel, where we classify centroids to their nearest angular neighbor.

For side classification comparisons, the two centroids corresponding to a given odor presented on both sides are removed. From the remaining 28 centroids, two points representing ipsilateral and contralateral cortex are made by averaging over all remaining centroids from a particular side. The two points we removed are then classified as ipsilateral or contralateral depending on which mean they are nearest to. This procedure is repeated 15 times, removing a different odor each time. We report the average accuracy and standard deviation over 200 randomly chosen pseudopopulations in Figures 4 and S5.

### Computational Modeling

We built a feed-forward neural network model to explore the implications of a random or structured connection between the olfactory cortex in each hemisphere. We effectively model one cortical region in each hemisphere, based on the AON.

To begin, we considered the cortical response to odors presented ipsilaterally, later we will link these to explore interhemispheric effects. Each cortical neuron’s activity to an ipsilateral odor o is a weighted sum of glomerular activities passed through an activation function:

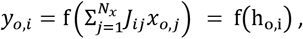

where **J** is the glomerular to cortical connectivity matrix, *x*_*o*,*j*_ *and y*_*o*,*i*_ are the firing rates of glomerulus j and cortical neuron i respectively to odor o, h_o,i_ is the input to cortical neuron i, j runs from 1 to *N*_*x*_ (the total number of glomeruli), and f is the activation function.

The zero point of these neurons is set to be basal firing rate plus response to a blank odor. Therefore, we modeled only odor related perturbations from the blank and base to match all comparisons to the blank and base subtracted data.

The nonlinearity, *f*, was chosen to recreate features of the electrophysiological recordings. First, it was found that the odor related perturbations are equally likely to be positive or negative (Of all the significant neuron-odor pairs, 46% are positive in the AON, 48% in the APC, and 48% in the PPC) therefore we chose a nonlinearity that was symmetric around zero. Negative activity does not mean the neurons have a negative firing rate, but that their firing rate is inhibited relative to their response to base plus blank. Second, consistent with previous findings (Bolding and Franks, 2017; Iurilli and Datta, 2017; Poo and Isaacson, 2009), the cortical response to odors was relatively sparse. Therefore, we chose the shrink function:

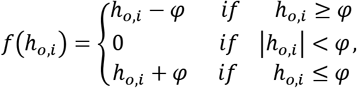

where ϕ is a threshold that controls sparseness.

Based on previous work we then modelled the connectivity matrix between the bulb and the cortex to be completely random (Schaffer et al., 2018) with elements either equal to 0, with probability 1 − ξ, or drawn independently from a zero mean distribution with finite variance 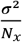, with probability ξ. ξ, therefore, controls the sparseness of the matrix^1^. Now by Central Limit Theorem ho,i is distributed in a Gaussian manner and with zero mean.

Since input to each cortical neuron is statistically independent, to fully specify its distribution, we must simply set the odor-odor covariance matrix of the Gaussian. Under the assumption that the size of the OB output response is identical for each odor (|***x***_***o***_|^**2**^ = |***x_o′_*** |^**2**^ ∀ odors o and o’), which can be justified by the divisive normalization that appears to happen in the OB (Banerjee et al., 2015; Zhu et al., 2013), the elements of the covariance matrix can be found as follows:

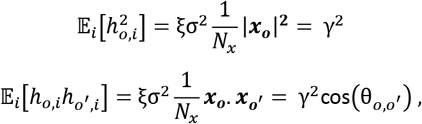

where 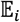 refers to an expectation over the cortical population and we defined a new variable 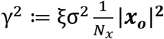 that controls the variance, and θ_*o,o′*_ is the angle between two glomerular representations. For the 15 odors under consideration, we used calcium imaging to extract the cosine similarity which serves as the Gaussian covariance. To complete our ipsilateral specification we must, therefore, only choose γ and a threshold ϕ – we shall rationalize these terms in two ways.

The threshold controls sparseness of cortical response, this we match to the AON recordings. The sparseness of the recordings is measured as the proportion of neuron-odor pairs for which the Wilcoxon signed rank test p value comparing the set of seven trials to mean blank odor response is below 5% (22.4% Ipsilateral, 25.2% Contralateral). Labeling the sparseness as S, because of the assumed Gaussianity of cortical inputs, we get:

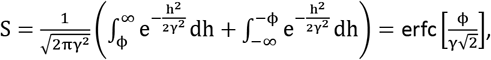

where erfc is the cumulative error function. Therefore, the sparseness sets the ratio of threshold to variance:

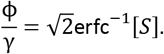

Finally, we use the ipsilateral distribution of significant response magnitudes in the AON (Figure 2E, top panel) to fit the conversion of numbers to firing rate (Hz), γ. Our model predicts the significant response magnitude distribution to be the tail of a Gaussian, by changing γ we are simply rescaling the x axis of this distribution, we choose the γ that permits the best model-data fit. This fully specified the ipsilateral simulations and fit the data remarkably well (see Figures 6 and S7).

To compare ipsilateral to contralateral responses we create two ipsilateral responses according to the scheme above. One of these is then mapped through a cross cortical connectivity matrix to create a contralateral response. We use this contralateral response and the second ipsilateral creation for our comparisons to mice.

Our final task, therefore, is to design a cross-cortical connectivity matrix. In the body of the paper, we use a simple Hebbian structured matching of each odor between hemispheres. Here we include two additional connectivity schemes that are low dimensional approximations to G_Struct_. The first uses SVD to create a low rank matrix closest to G_Struct_ as measured by the Frobenius norm. The second uses the principal components to create G_Struct_ using a smaller number of Hebbian matching experiences. These show that the particular form of the structured connectivity is not important. In each case we combine the structured matrix with a random matrix and examine how much structure is required to fit the observed data.

The random component has elements drawn from a zero mean normal distribution with variance 1 (this choice of variance is unimportant as the matrix will be rescaled in future):

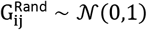

with probability ξ, else 0.

And the structured Hebbian component maps from contralateral to ipsilateral odors (Dayan and Abbott, 2001):

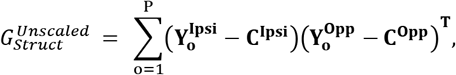

where 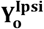 is the response vector to odor o presented ipsilaterally in the ipsilateral cortex, **C**^**Ipsi**^ is response in the ipsilateral cortex averaged over odors, P=15 is the total number of odors, and 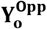 is the response in the opposite cortex to an odor presented in the opposite nostril.

These two matrices currently have outputs that are of arbitrary magnitudes. To ensure each contribution has a roughly similar sized output, the structured section is rescaled by a factor:

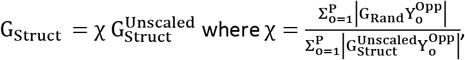

before being combined using the parameter α to create a cross cortical link with varying levels of randomness/structure:

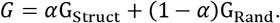

Since the size of the outputs of each matrix are equal, we can now interpret alpha as the proportion of contralateral input magnitude that is formed by structured connectivity.

The magnitude of the elements of this matrix are then scaled again such that the resulting contralateral representation has the desired sparseness when the same piriform threshold is applied as was used to create the ipsilateral sparseness. Finally, the conversion to Hz, γ, is used to compare to data.

We varied alpha and looked at how four measures of cross-cortical alignment changed (Figures 6E-6J; see main text for details) and how simulations compared to our actual AON data. We chose the optimal alpha to be the point with the minimum z^2^-score across the four – the smallest squared sum of sigmas (error bars) between the point and the AON data line in all four plots.

We modeled our AON with 50,000 neurons, this is a rough underestimate based on other paleocortical neuronal densities (Srinivasan and Stevens, 2017) and measurements of AON size from the Allen Brain Atlas. Our model, however, was robust to changes in number of neurons, for example we tried a 25,000-neuron version (Figures S7E-S7H) and the optimal alpha did not change.

Our first low rank approximation uses Singular Value Decomposition to factorise G_Struct_ :

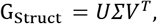

where U and V are orthogonal matrices and *Σ* is a diagonal matrix. We vary the dimensionality of our approximation through setting all but the largest Q singular values in *Σ* to 0, creating *Σ*\* which we then use to recreate G_Struct_ :

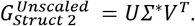

The last type of structured connectivity is a simple extension, comprising a Hebbian matching of vectors between the two hemispheres. However rather than matching the odors explicitly we try to align the odorant subspace in which the odorants sit. To do this we calculate the principal components of the contralateral representation and design a connectivity scheme that aligns the first few of the contralateral principal components between cortices:

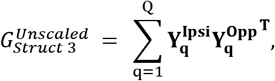

Where Q is the number of principal components used, 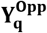 is the q-th principal component of the opposite cortical representation, and 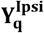 is the ipsilateral representation of the same mixture of odors that comprises 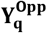.More explicitly, the q-th principal component in the opposite cortex is a mixture of normalized odorant vectors:

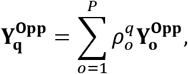

Where ρ^q^ is the vector of coefficients for the q-th principal components. We then use the same set of coefficients to create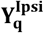:

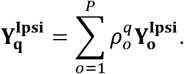

The dimensionality of each approximation, Q, must be chosen such that most of the response variation is captured; however, the low dimensionality of the responses means high alignment can be achieved despite a small Q. For example, in Figures S8A and S8B we show how cross-cortical decoding accuracy varies with the dimensionality of each approximate structured matrix. Only 7 singular values (Figure S8C-S8F), or 6 principal components (explaining 73% of the variance, Figures S8G-S8J), are needed for the alignment metrics to reach the values seen in neuronal recordings.

### Quantification and Statistical Analysis

All quantifications and statistical analyses were performed using custom scripts in MATLAB (MathWorks), except (1) the circular statistical analyses, which were performed with the CircStat toolbox written by Philipp Berens (Berens, 2009), and (2) the Hartigan’s dip tests, which were performed with the code written by Ferenc Mechler (Mechler and Ringach, 2002). The linear regressions were obtained with the least square method. The data is given as median ± standard deviation (SD), unless specified otherwise. For all statistical tests, the critical value was set to 5%. All statistical tests were two-sided, with the exception of the bootstrap strategies (where we asked whether the measurements were significantly higher than chance).

Additionally, the significance of the linear regressions was tested using either an F-test (Figures 3, S3, and S4). The significance of the percentage of bilaterally correlated neurons was tested using a bootstrap strategy (see figure caption for details). Table S2 summarizes all the statistical tests performed in this study

## Data and Code Availability

All data and code used to compute quantities presented in this study is available on the Murthy lab GitHub page: https://github.com/VNMurthyLab/IpsiContra_v2.

## Author Contributions

V.N.M and J.G designed the experimental setup. C.P and W.D. designed the computational modeling strategy with inputs from all authors. J.G. acquired the electrophysiological data. J.G. and W.D. analyzed the data. J.G. and W.D. wrote the manuscript with inputs form all authors.

## Acknowledgements

Research reported in this paper was supported, in part, by grants from the NIH (R01 DC016289, R01DC011291), the Harvard Merit Fellowship, and the Herchel Smith Scholarship. We thank Vikrant Kapoor, as well as all members of the Murthy and Pehlevan labs for helpful feedback. We also thank Massimo Vergassola and Gautam Reddy for critical discussion. Finally, we thank Naoshige Uchida and Kenneth Blum for their important feedback on a previous version of this manuscript.

## Supplementary Information

### Note S1: Random Connections Produce Zero Correlations – a Derivation

We will prove that if the same response is mapped through two different random matrices the resulting representations will be uncorrelated. Let **x** be our *N*-dimensional input pattern, for example the bulb response to an odor. Project it through two different random matrices to the same set of cortical neurons. The two matrices, **J** and **G**, each have independently identically distributed elements drawn from a mean zero distribution with finite variance. This creates two further representations which we shall later pass through an activation function. First, however, we shall consider the pre-thresholded representations:

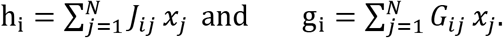

We can use the Multivariate Central Limit Theorem to argue that these two representations are jointly normally distributed with zero mean:

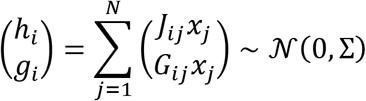

The correlation matrix can be derived and shown to be diagonal:

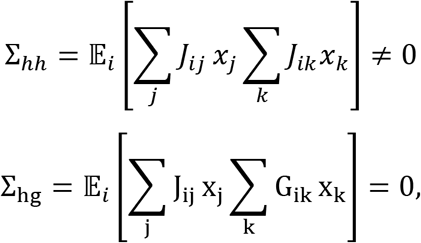

where the last equality follows from the independent, zero mean, nature of elements of **G** and **J**.

Now, since h_i_ and g_i_ are jointly normally distributed random variables with a diagonal covariance matrix, they are also independent. We then pass these representations through a nonlinearity. Using the fact that functions of independent variables are independent, the resulting output representations are also independent and hence uncorrelated.

This demonstration can be extended to many cases of direct interest. If we let *h* and *g* depend on different odors they are still independent and uncorrelated by a simple generalization of the previous argument:

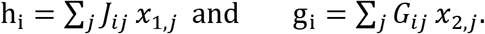

The experiments consider another similar case. Interpret h (g) as the unthresholded representation in the ipsilateral (contralateral) AON to an odor presented ipsilaterally (contralaterally). If the cross cortical matrix is random then thresholding these responses and mapping them cross cortically will not change their independence from one and other.

Previous work has shown that, if two correlated representations are projected through a random matrix, the resulting representations remain correlated (Babadi and Sompolinsky, 2014; Schaffer et al., 2018). The key difference in our work is that we are considering two different random matrices, one in each hemisphere. Hence, this derivation has shown that projecting a pair of correlated representations through two different random matrices eliminates the correlations between resulting outputs.

Therefore, since observed correlations between odors presented ipsilaterally and contralaterally are not zero, the cross cortical connectivity must be structured in some way.

**Figure S1:**
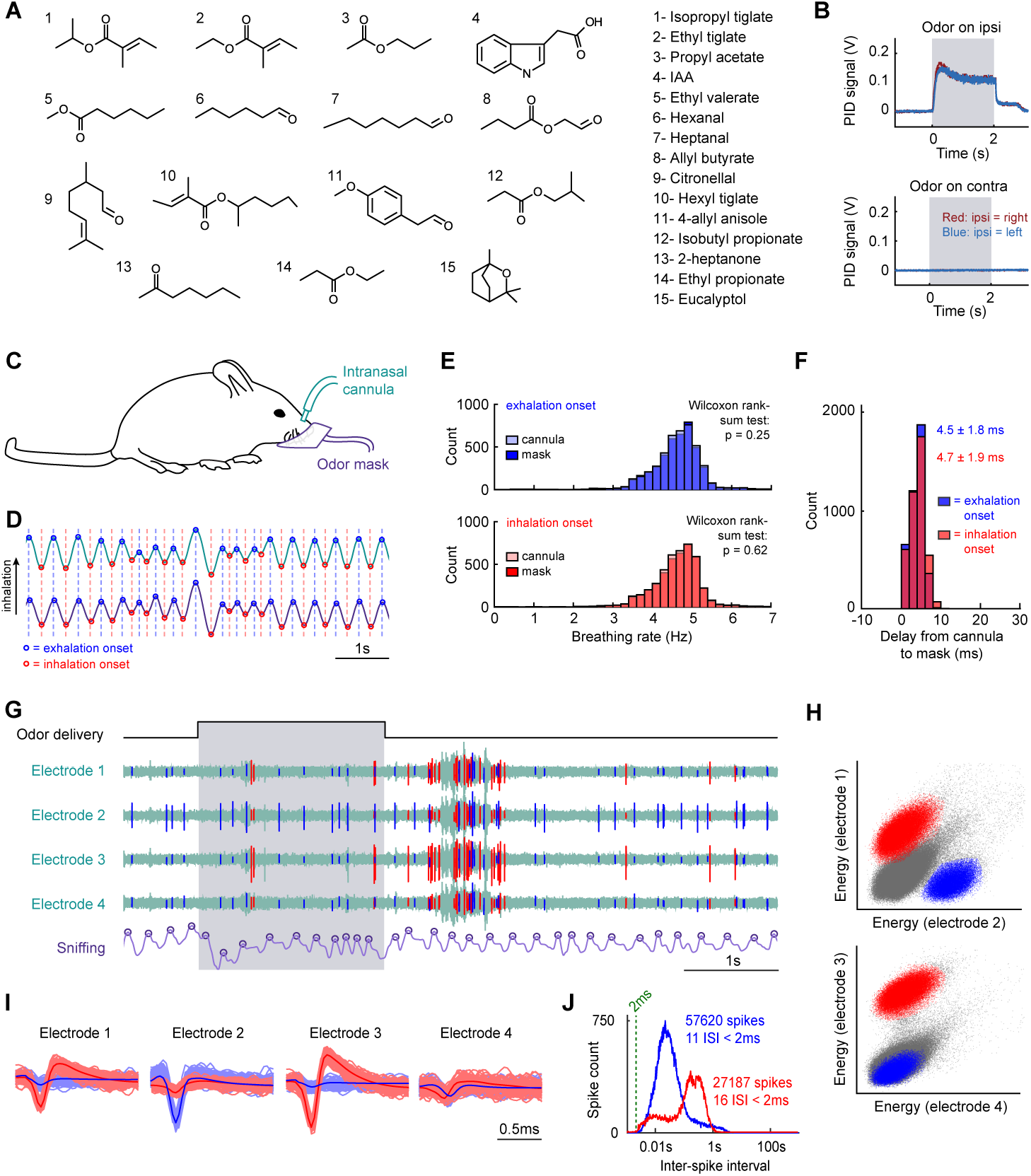
A New Method for Unilateral Odor Delivery. ***Related to Figure 1*. (A)** Odorants used in this study. **(B)** Control of the symmetry of the olfactometer. A PID measured the changes of odor concentration over time at the end of the right or the left end of the facial mask. Each measurement was repeated three times; therefore, each graph shows six over-imposed traces, three with the PID on the right, three on the left. Note the similarity between the traces obtained on the left and right of the mask, and the absence of odor detected on the contralateral side. Grey: odor presentation (here: isopropyl tiglate). All 15 odors were tested and showed similar results. **(C)** Diagram of the setup used for assessing the reliability of the face mask for respiration monitoring. The breathing of awake mice was monitored simultaneously with the mask and a pressure sensor connected to a chronic intranasal cannula (n = 3 animals, 15 min of continuous recording per mouse). **(D)** Exemplar breathing traces, simultaneously recorded from the intranasal cannula (green) and the face mask (purple). Note the synchrony of both signals and the apparent absence of phase shift. **(E)** Histogram of instantaneous breathing rates, calculated with the exhalation (top panel) or inhalation (bottom panel) onsets. The data shown here comes from the full monitoring session of one exemplar mouse. The distributions obtained from the intranasal cannula (light bars) or the face mask (dark bars) are almost identical. **(F)** Delay from cannula to mask, calculated with the exhalation (blue) or inhalation (red) onsets. The data from the three mice were combined here. Values: median ± SD. The phase shift between the intranasal cannula and the face mask is small and reliable. **(G)** Example of tetrode recording. The four electrodes belong to the same tetrode. Grey: odor delivery. Green: unsorted electrode traces. Blue and red: two single units. Purple: sniffing acquired simultaneously (up: inhalation). These exemplar traces were recorded in mouse APC1. **(H)** Example of single unit clustering. The red and blue units are the same as in (G). **(I)** Example of clustered single units. The red and blue units are the same as in (G). Lighter traces: all the peaks from the recording session were over-imposed. Darker traces: average peaks. **(J)** Spike count versus inter-spike interval (ISI) for the two units shown in (G) (blue and red). Note the presence of a refractory period.

**Figure S2:**
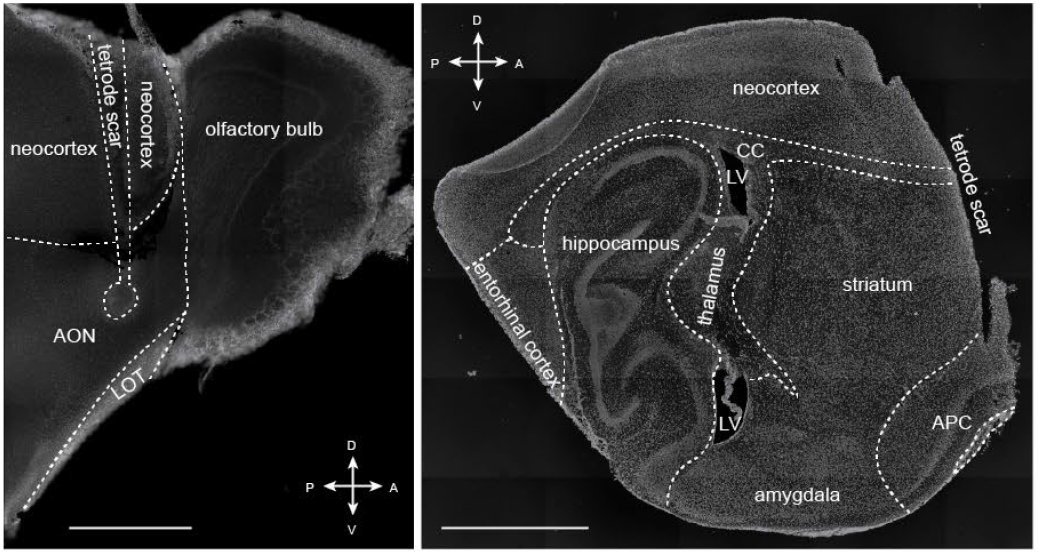
Post-Mortem Confirmation of Tetrode Placement in the Olfactory Cortex. ***Related to Figure 2*.** Sagittal sections of two exemplar animals, implanted in the AON (left) and APC (right). The round shape on the ventral end of the tetrode scar in the left panel was caused by post-mortem electrolesion. For animals recorded in the APC, the tetrodes went all the way through the brain, until they reached the ventral part of the skull. As a consequence, it was not possible to save the most anterior part of the section in the right panel. A: anterior. P: posterior. D: dorsal. V: ventral. CC: corpus callosum. LV: lateral ventricle. LOT: lateral olfactory tract. Scale bar: 0.25mm (left) or 1mm (center, right). DAPI staining. We did not trace the border between striatum and amygdala as we could not confidently determine it.

**Figure S3:**
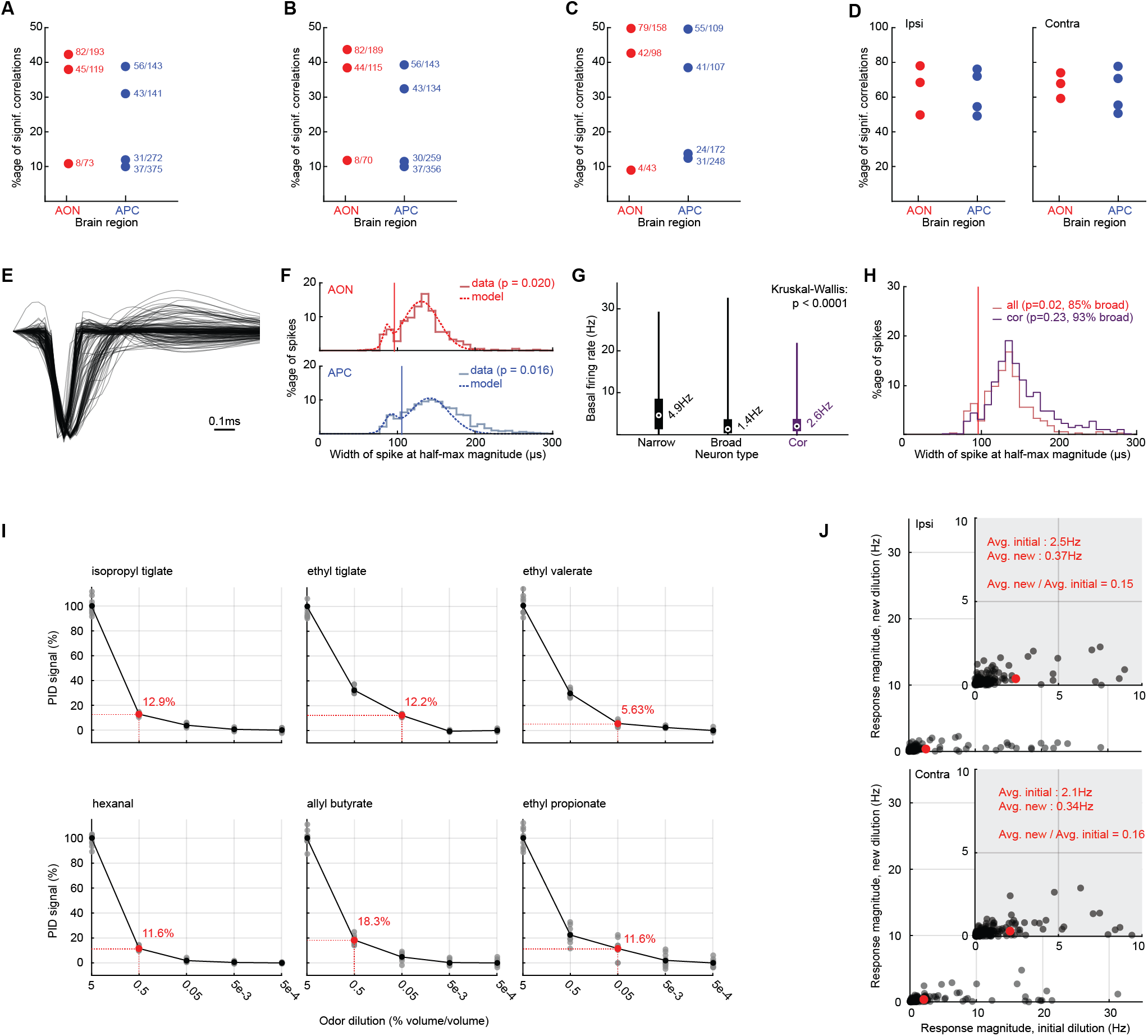
Bilaterally-Correlated Neurons. ***Related to Figure 3*. (A) to (D)** Percentage of bilaterally-correlated neurons per mouse. **(A)** Identical to Figure 3B. **(B)** Same as Figure 3B, except we kept only the neurons responding to at least one odor on one side for this analysis. **(C)** Same as Figure 3B, except we kept only the neurons responding to at least one odor on each side for this analysis (note that the odors eliciting responses can be different on each side). **(D)** Correlation across artificially-generated trials. For each mouse, each side, each neuron, each odor, two sets of 7 artificial trials were generated using a Poisson process. Then for each side, an analysis similar to Figure 3B was performed, except we looked for correlations between the first and second sets of 7 artificial trials, instead of correlations between ipsi and contralateral trials. A full circle means the percentage is higher than chance (significance is determined with a bootstrapping method similar to Figure 3C). Panel (D) allows us to estimate the apparent percentage of bilaterally-correlated neurons one would expect if all neurons were bilaterally-correlated. **(E)** Spike traces from mouse PPC1. The average spike of all the neurons recorded from APC1 are shown. The spike amplitudes were normalized for display. Note the apparent bimodal distribution of spike widths. **(F)** Histograms of the spike widths at half-maximum amplitude, per OC region. For each histogram, a Hartigan’s dip test was performed to confirm that the distribution of spike widths was not unimodal (AON p = 0.020, APC p = 0.016). The corresponding p-values are reported next to each graph. We also used a maximum likelihood estimate approach to fit a mixture of two Gaussian distributions to each distributions (AON: weight of the 1st Gaussian relative to the mixture w = 0.07, μ1 = 85.6, σ1 = 4.5, μ2 = 130.1, σ2 = 22.9; APC: w = 0.11, μ1 = 94.2, σ1 = 8.1, μ2 = 146.3, σ2 = 26.2). For each distribution, the vertical bar indicates spike width at which the Gaussian mixture reaches its minimum between the two peaks: we chose this value as the definition for “narrow” and “broad” spikes (AON 95μs; APC 110μs). Using this criterion, we found 85% of broad spikes in the AON and 89% in the APC. **(G)** Basal firing rate of the neurons with narrow versus broad spikes. Based on the histograms in (F), spikes were categorized as narrow or broad. White circle and value next to it: median. Thick bar: quartiles. Thin bar: range of values. The neurons with narrow spikes showed a significantly higher basal activity than the broad ones. This finding, along with the presence of a bimodal spike width distribution, suggests that the neurons with narrow spikes are mostly inhibitory, while the neurons with larger spikes are mostly excitatory. The purple bar shows the same analysis performed on the bilaterally-correlated (noted “cor”) neurons of the AON and APC (Kruskal-Wallis test across the “narrow”, “broad’, and “cor” categories: p = 3.2e-6; post-hoc Wilcoxon rank-sum test, “narrow” versus “broad” p < 0.0001, “narrow” versus “cor” p = 0.0008, “broad” versus “cor” p = 0.60). **(H)** Comparison of the spike width distribution of all AON and APC neurons versus bilaterally-correlated neurons. Unlike the overall distribution in the AON and APC, the spike width distribution for the bilaterally-correlated neurons is not significantly different from a unimodal distribution (Hartigan’s dip test, p = 0.06) Almost all the bilaterally-correlated neurons have a broad spike (97% neurons have a broad spike), suggesting they are putative excitatory neurons. **(I)** PID signal amplitude at different odor dilutions. All amplitudes are reported relative to the amplitude of the dilution used for the rest of this study (5%). Each grey dot is one PID measurement (10 repeats per condition), the black dots are the average amplitudes. For each odor, the red dot indicates the dilutions at which the PID signal is about 10 times weaker than the initial 5% dilution. **(J)** Response magnitude of the initial versus weaker dilutions. Each graph is one side of the mask. Each dot is one neuron-odor pair (n = 2 mice recorded in the AON, 120 neurons). We only tested the 6 odors shown in panel I (initial dilution: 5%; new dilution: see red dot in I). Only the neuron-odor pairs significantly responding to the initial 5% concentration are shown. Red: average for each side. When we stimulated the same side of the mask with 10 times less odorant, we obtained neuronal responses about 6 to 7 times weaker than with our initial 5% concentration. Therefore, cross-contamination in the mask, if any, cannot explain the presence of side-invariant odor responses.

**Figure S4:**
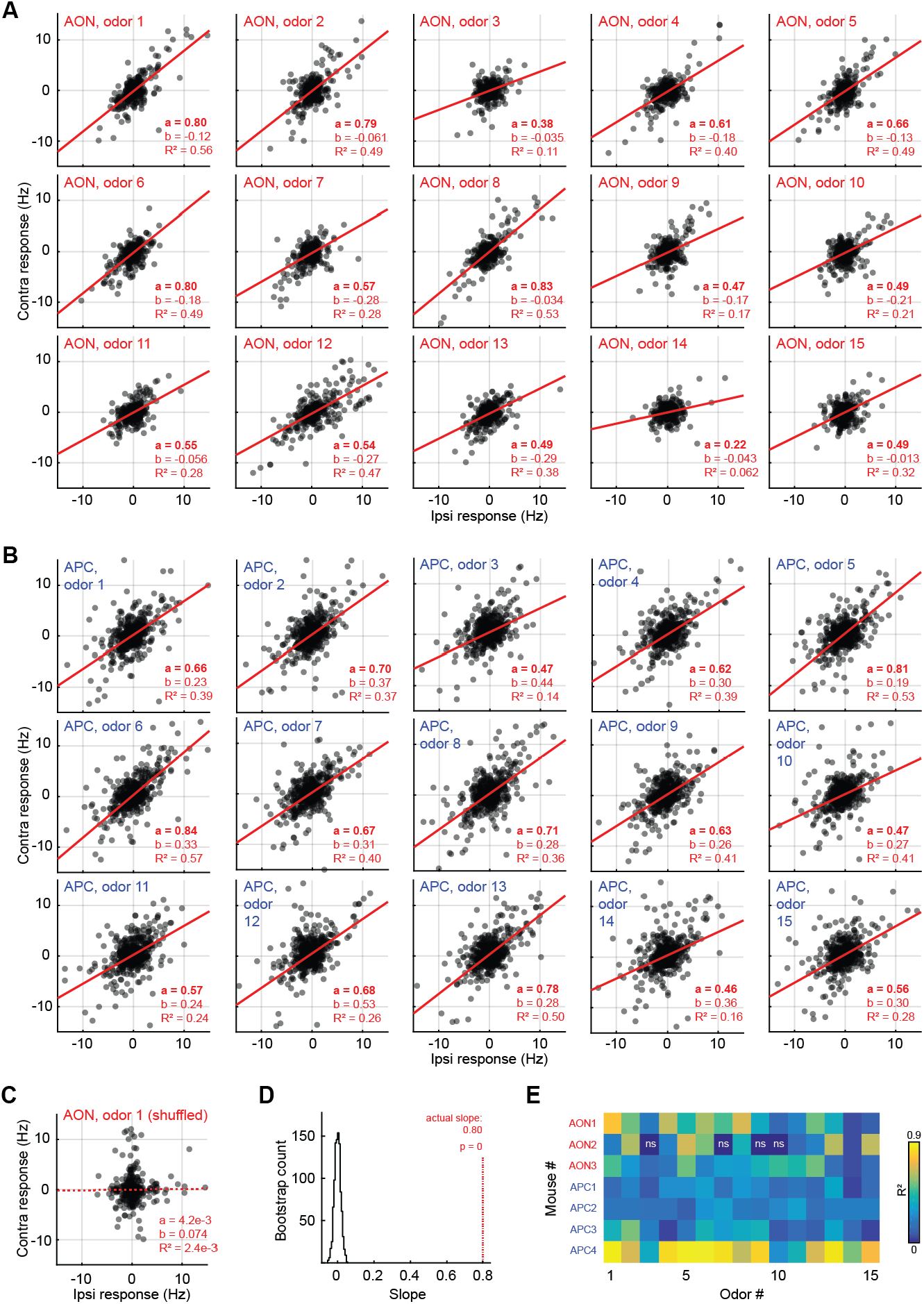
Correlation of Population Odor Representations. ***Related to Figure 3*. (A)** and **(B)** Ipsi- versus contralateral responses per odor in both OC regions tested. (A) AON. (B) APC. Red line: linear regression (contra = a × ipsi + b). Solid lines show significant regressions (F-test, p < 5%). Odor # is the same as Figure S1A. Each dot is one neuron. The top left graph for each region is also shown in Figure 3G. **(C)** Same as panel (A), odor 1, except neuron identity has been randomly shuffled on each side. Note the absence of significant correlation. **(D)** Significance of the regression slope for panel (A), odor 1. We built a chance distribution by repeating the shuffling shown in panel (C) 1000 times. We then compared the actual value of the slope to this distribution to determine its significance. Here we show the distribution of chance slopes for the AON, odor 1. The results of our bootstrap strategy matched the results of the F-test mentioned above. **(E)** We performed the same analysis as panels (A) and (B), but we looked at each mouse separately. Here we report the R^2^ value of each regression, as well as whether the correlation was significant (F-test). N.s. means that the correlation was not statistically significant (p ≥ 5%).

**Figure S5:**
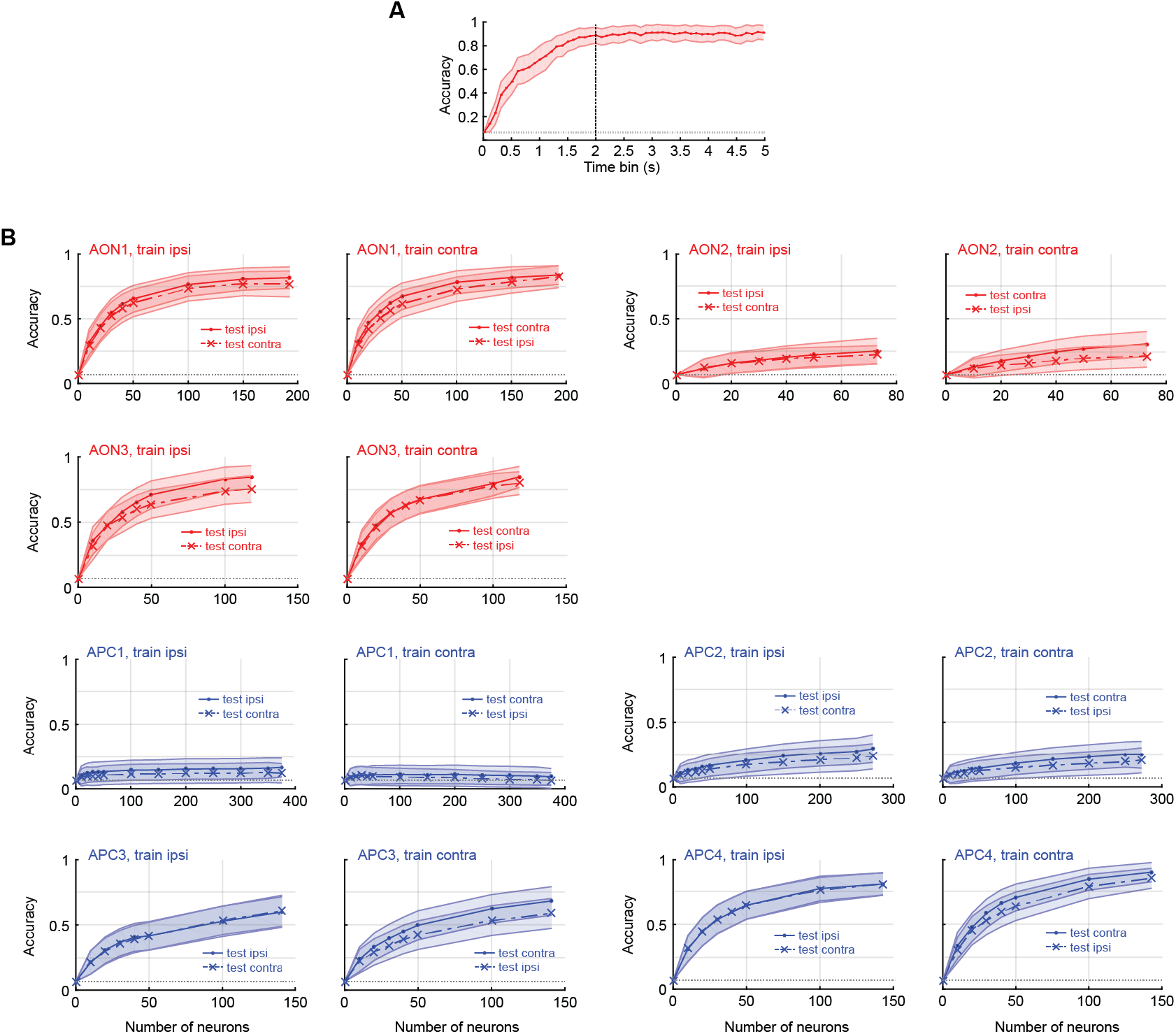
Ipsi- and Contralateral Odor Decoding. ***Related to Figure 4*. (A)** Decoding accuracy with varying odor response time windows. For each data point, response vectors were built by summing spikes from 0s to the time indicated on the x-axis (responses aligned to the first sniff after odor onset, see Methods for details). This example shows the odor decoding for all neurons in the AON, train ipsi, test ipsi. Odor decoding accuracy is maximal at 2s, therefore we decided to perform all our decoding analysis over an odor response window of 2s. **(B)** Same as Figures 4B-4C, except the decoding process was applied to each mouse separately. Accuracy at n = 50 neurons is significantly higher than chance for all mice. The mice with poorer decoding accuracy were usually the ones in which neurons showed fewer / weaker responses on average, as well as fewer bilaterally-correlated neurons (see Figure S6).

**Figure S6:**
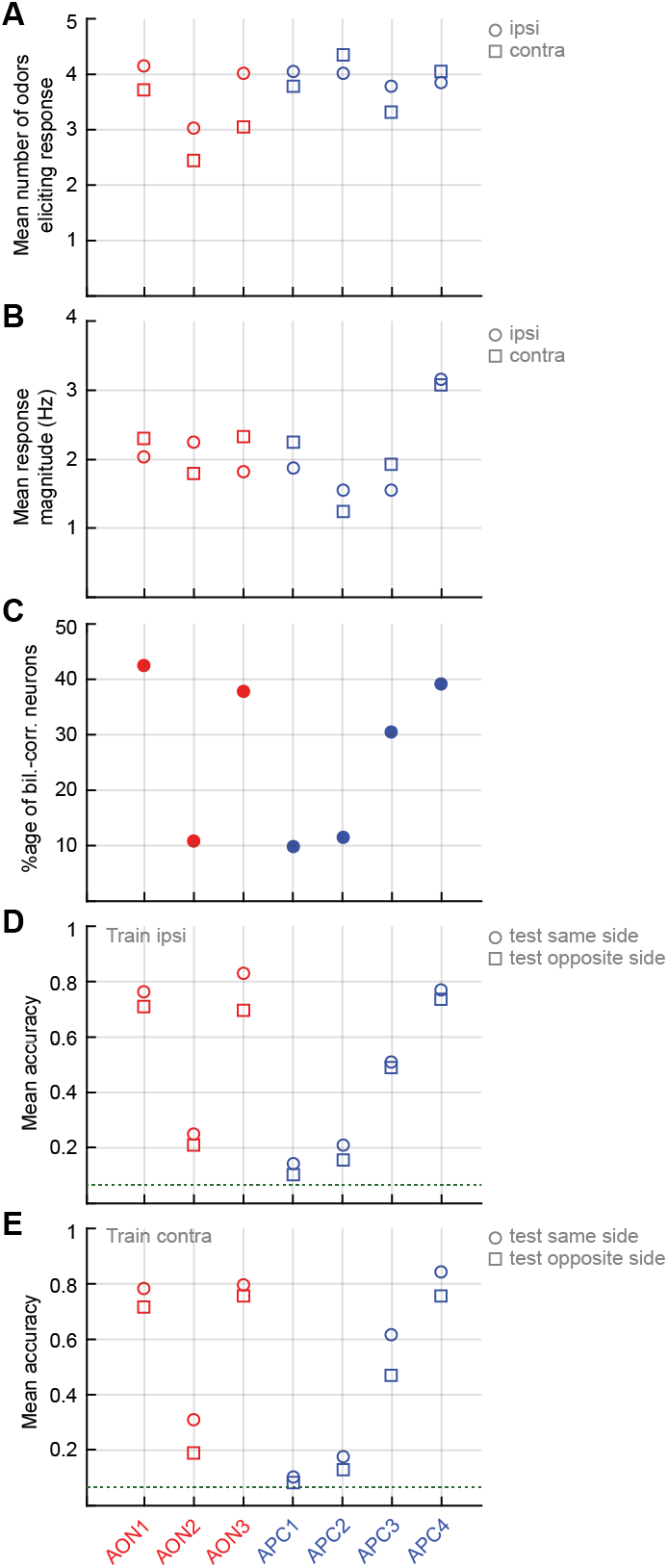
Variability Across Mice. ***Related to Figures 3 and 4*.** In this figure we point at factors that may explain the variability we observed across mice in terms of percentage of bilaterally-correlated neurons, as well as odor decoding accuracy. **(A)** Mean number of odors eliciting response (related to Figure 2E). **(B)** Mean response magnitude (related to Figure 2F). **(C)** Percentage of bilaterally-correlated neurons (related to Figure 3B). Solid dot: significant percentage. **(D)** Mean accuracy of the odor decoding process with model trained on responses to ipsilateral presentations (related to Figure S5B). **(E)** Similar to (D) with training on contralateral presentations.

**Figure S7:**
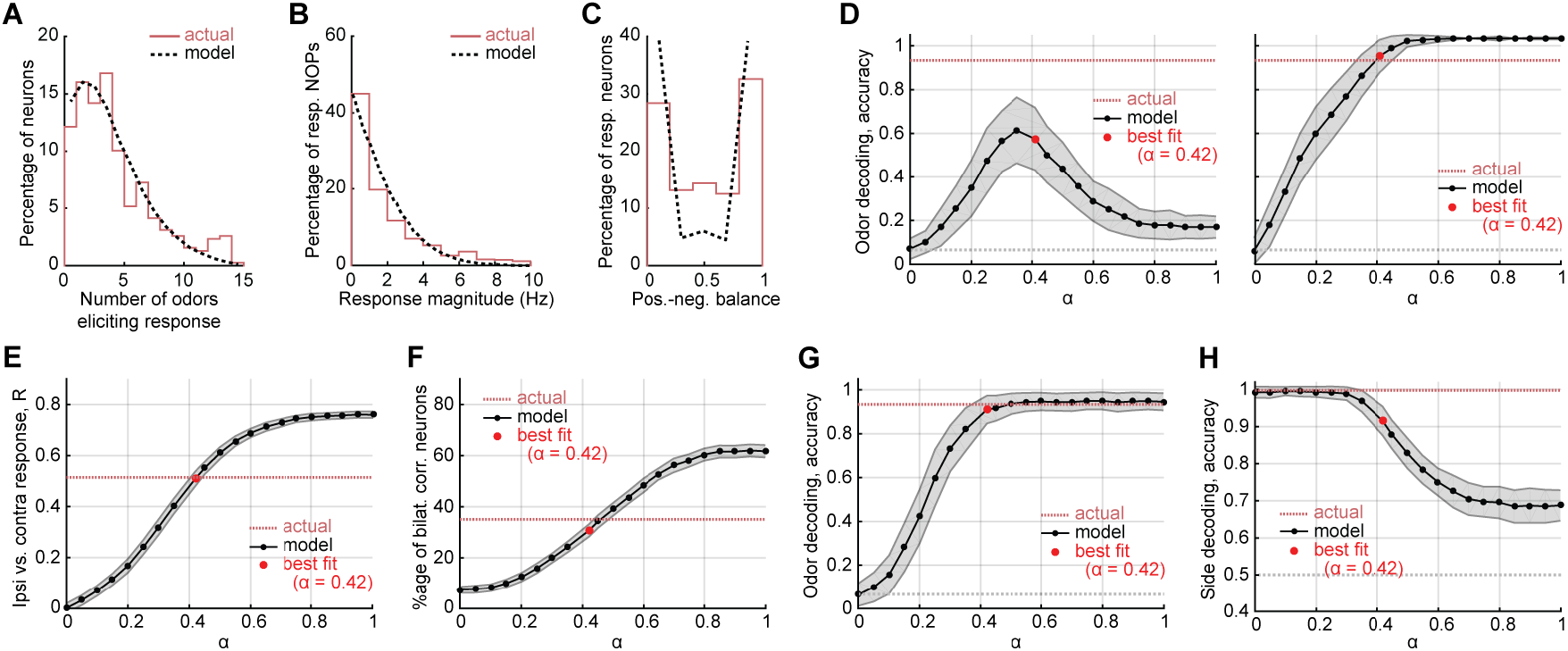
Modeling Olfactory Cross Cortical Connections. ***Related to Figure 6*. (A) to (C)** Same as Figures 6B-6D, except for we are comparing the distributions of mouse ipsilateral data and our simulations of ipsilateral responses with optimal alpha (0.42). Mouse data is identical to the contralateral lines from Figures 2E-2F, top panels. **(D)** Same as Figure 6I, except we decoded ipsilateral odors using the contralateral presentations. (Left) Using closest Euclidean neighbor decoding is markedly worse in simulations than mice, due mainly to the difference in magnitude of responses that results as you increase the structure parameter. (Right) Using closest angular neighbors instead recaptures better performance, and the best alpha overlaps with the AON performance. This suggests that, in the left panel, the decoder limited performance rather than lack of information. **(E) to (H)** Same as Figures 6F, 6G, 6I, and 6J respectively, except each OC layer contains 25,000 neurons (instead of 50,000 neurons in Figure 6, see Methods). These show that the optimal alpha level (0.42) is robust to changes in the number of cortical neurons simulated.

**Figure S8:**
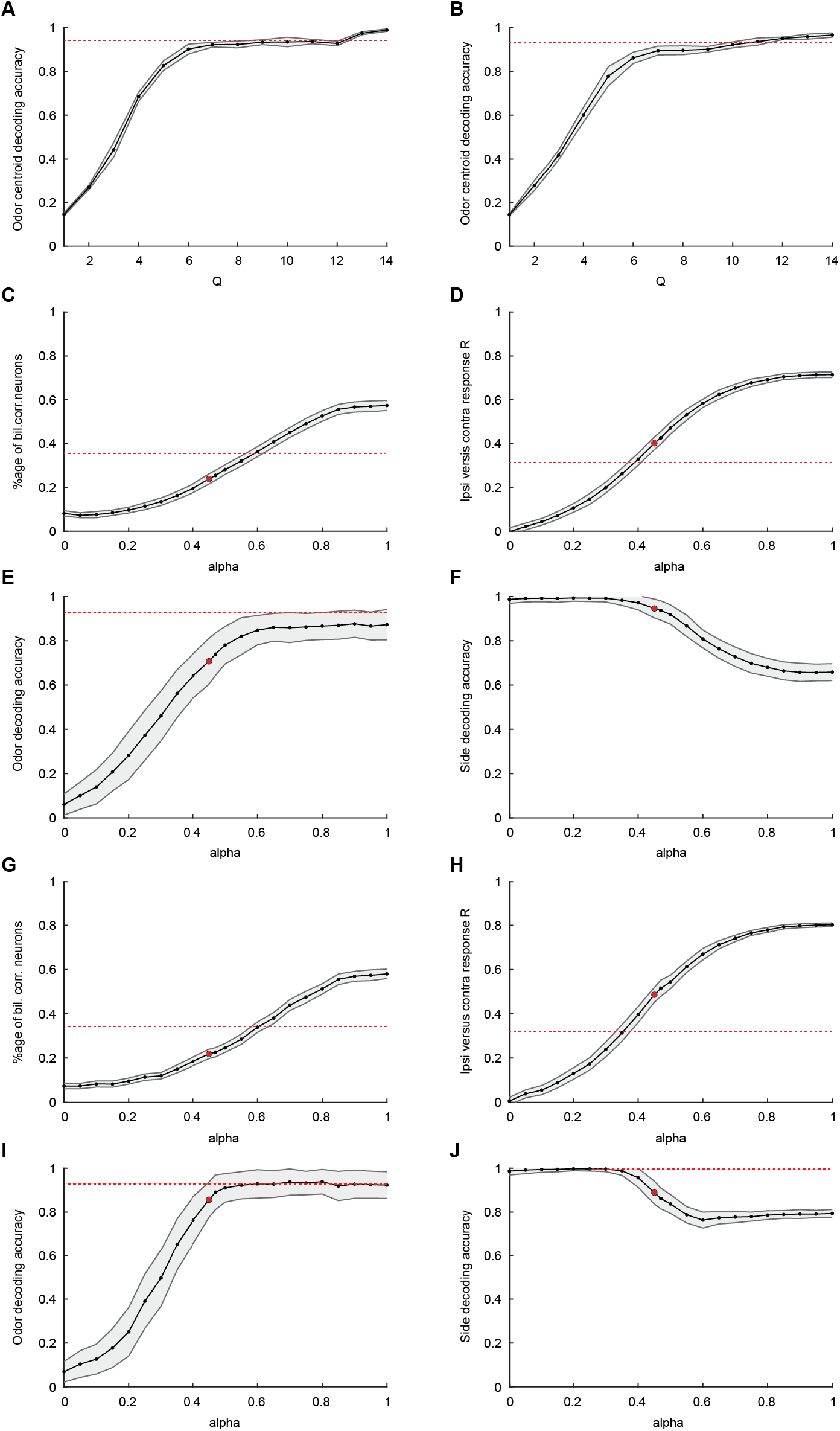
Low-dimensional structured connectivity is sufficient to produce alignment. **(A)** and **(B)** Odor decoding accuracy using 385 neurons as Q is varied, averaged over 200 iterations of decoding and 10 different creations of *G_Struct_*, **(A)** approximates using SVD and **(B)** using PCA. **(C) – (F)** are the same as Figure 6F, G, I, and J but using an approximate *G_Struct_* created with 7 singular values. **(G) – (J)** are the same but using 6 PCA components.

**Table S1:** Mice Recorded in this Study. All tetrode recordings were performed in awake, head-restrained mice.

| Brain region | Mouse name | Age at the first recording | Number of sessions | Number of neurons |
| --- | --- | --- | --- | --- |
| AON | AON1 | 3 m.o. | 11 | 193 |
|  | AON2 | 3 m.o. | 17 | 73 |
|  | AON3 | 3 m.o. | 14 | 118 |
| APC | APC1 | 3 m.o. | 15 | 375 |
|  | APC2 | 4 m.o. | 12 | 272 |
|  | APC3 | 3 m.o. | 11 | 141 |
|  | APC4 | 3 m.o. | 11 | 143 |
| PPC | PPC1 | 5 m.o. | 12 | 184 |
|  | PPC2 | 5 m.o. | 11 | 155 |
|  | PPC3 | 4 m.o. | 10 | 166 |
| OB | OB1 | 7 m.o. | 1 | 7 |
|  | OB2 | 7 m.o. | 1 | 9 |
|  | OB3 | 8 m.o. | 1 | 11 |
|  | OB4 | 8 m.o. | 2 | 15 |

**Table S2:**
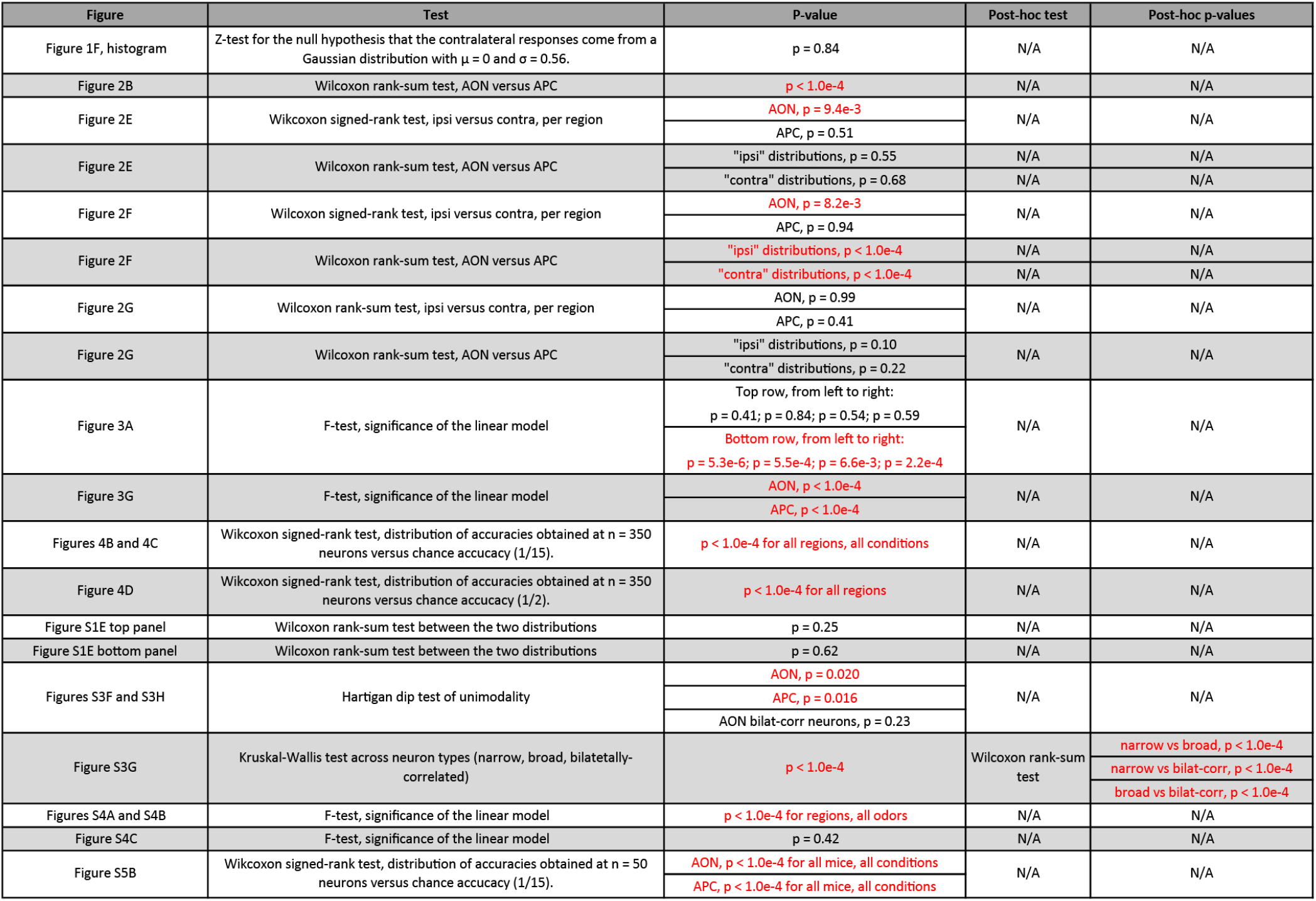
Statistical Tests Performed in this Study. The statistical tests showing significant differences are in red (critical value: 5%). For post-hoc tests, the p-values reported in the table have been corrected for multiple testing problem (Bonferroni method).

## Footnotes

1 Note that negative weights do not mean that OB inputs are directly inhibitory; they rather model OB inputs projecting to inhibitory OC neurons, which in turn project to our OC neurons of interest.

